# Sub-3 Å resolution protein structure determination by single-particle cryo-EM at 100 keV

**DOI:** 10.1101/2024.09.05.611417

**Authors:** Dimple Karia, Adrian F. Koh, Wen Yang, Victoria I. Cushing, Benjamin Basanta, Daniel B. Mihaylov, Sagar Khavnekar, Ondřej Vyroubal, Miloš Malínský, Ondřej Sháněl, Vojtěch Doležal, Juergen M. Plitzko, Lingbo Yu, Gabriel C. Lander, A. Radu Aricescu, Basil J. Greber, Abhay Kotecha

**Affiliations:** Materials and Structural Analysis Division, Thermo Fisher Scientific, Achtseweg Noord 5, 5651 Eindhoven, NL; The Institute of Cancer Research, Chester Beatty Laboratories, 237 Fulham Road, London SW3 6JB, UK; Dept. of Integrative Structural and Computational Biology, Scripps Research, La Jolla, CA 92024, USA; MRC Laboratory of Molecular Biology, Francis Crick Avenue, Cambridge, CB2 0QH, UK; Max Planck Institute of Biochemistry, CryoEM Technology, Martinsried, Germany; Present address: Arzeda, Seattle, WA, USA; Present address: Thermo Fisher Scientific, Achtseweg Noord 5651 Eindhoven, NL

## Abstract

Cryo-electron microscopy (cryo-EM) has revolutionized structural biology by providing high-resolution insights into biological macromolecules. Here, we present sub-3 Å resolution structures determined using the 100 keV Tundra cryogenic transmission electron microscope (cryo-TEM), equipped with the newly developed Falcon C direct electron detector (DED). Our results demonstrate that this lower voltage microscope, when combined with advanced electron optics and detectors, can achieve high-resolution reconstructions that were previously only attainable with higher voltage systems. The implementation of an extreme-brightness field emission gun (XFEG) and an SP-TWIN objective lens significantly enhanced the spatial and temporal coherence of the system. Furthermore, the semi-automated sample loader minimized contamination and drift, allowing extended data collection sessions without manual intervention. The high detective quantum efficiency (DQE) of Falcon C further improved the signal-to-noise ratio, which is critical for achieving high-resolution structures. We validated the performance of this microscope by determining the structures of various biological samples, including apoferritin, T20S proteasome, GABA_A_ receptor, haemoglobin, and human transthyretin ranging in size from 440 kDa to 50 kDa. The highest resolutions achieved were 2.1 Å for apoferritin, 2.7 Å for the 20S proteasome, 2.8 Å for the GABAA receptor, 5.0 Å for haemoglobin, and 3.5 Å for transthyretin. We also explored a larger specimen, a 3.9 MDa Adeno-associated virus (AAV9) capsid and resolved it a 2.8 Å. This work highlights the potential of 100 keV TEMs to make high-resolution cryo-EM more accessible to the structural biology community. Furthermore, it sets a precedent for the use of lower voltage TEMs in routine cryo-EM studies, not only for screening grids for single particle analysis but also for achieving high-resolution structures of protein samples.

## Introduction

Knowledge of three-dimensional structure is critical for understanding the molecular architecture, function, and regulation of protein complexes involved in key cellular processes. Over the past decade, cryogenic electron microscopy (cryo-EM) has become the predominant method for 3D structure determination of a wide variety of biological macromolecules and their complexes. Facilitated by significant advances in technology, which include the introduction of stable transmission electron microscopes (TEMs), high-speed direct electron detectors (DEDs) with improved detective quantum efficiency (DQE), and software increasing ease of data collection and processing and post-imaging drift correction when combined with the new detector advances, structural analysis has become straightforward and routine even for many challenging biological specimens (Kühlbrandt, 2014; Liao et al., 2013; Guaita et al., 2022).

The vast majority of novel experimental structures of biological macromolecules are now determined by single-particle cryo-EM on high-end 300 kV TEMs. While these microscopes have been the standard for delivering the highest resolution structures of proteins, nucleic acids, and their complexes, recent publications have shown that 200 kV cryo-TEMs with improved technologies, such as energy filters and DEDs, are also capable of producing high-resolution structures (Wu et al., 2020; Koh et al., 2022; Thangaratnarajah et al., 2022; Hamdi et al., 2020; Herzik et al., 2019). However, the costs of ownership and operation of high-end 300 kV and 200 kV TEMs present a considerable hurdle to adoption by new users (Vinothkumar and Henderson, 2016). 100 kV cryo-TEMs have been previously presented as a potential solution to achieve high resolution and reach a larger user base due to the reduced cost of microscope ownership and operation (Naydenova et al., 2019). Previous investigations suggested that there is a theoretical potential improvement of up to 25% in the amount of information available per unit damage for typical single particle cryo-EM specimens imaged using 100 keV electrons compared to 300 keV electrons (Peet et al., 2019). However, until recently, only hybrid detectors were demonstrated at 100 keV (Naydenova et al., 2019; McMullan et al., 2023), and there were no high-efficiency imaging DEDs with more than 1 million pixels and with software integration available that could satisfy the high imaging productivity requirements of single particle cryo-EM experiments. Additionally, as sample preparation and screening remain a hurdle for single particle analysis experiments due to the time required for optimization, routine access to cheaper instrumentation that can inform on sample quality could increase the efficacy of cryo-EM centers worldwide.

In this article, we present high-resolution structures reconstructed from images acquired on the 100 kV Tundra cryo-TEM with the Ceta-F detector and the newly developed Falcon C DED. We present results from both traditional long-duration (> 20 hours) data collections and quick screening runs (*∼*4 hours) and discuss the future potential of using 100 keV TEMs for single particle cryo-EM analysis.

## Results

In order to achieve a resolution of better than 3 Å using a 100 kV cryo-TEM, it is necessary to re-design the electron source and optics for optimal imaging performance. Traditionally, TEMs that operate at lower accelerating voltages (*≤* 120 keV), have been equipped with either a LaB_6_ or a tungsten filament. However, these electron sources have limitations such as low brightness (resulting in low spatial coherence) and large energy spread (resulting in low temporal coherence). These limitations cause the signal to drop below 50% at 5 Å resolution under commonly used low-dose imaging conditions (with dose rate of 10 e^-^/Å^2^/s, illumination area < 2 µm and total dose < 100 e^-^/Å^2^). To overcome this, a Tundra has been equipped with a 100 keV extreme brightness field emission gun (XFEG) (Figure 1a). The ultra-high brightness of an XFEG significantly reduces the effects of spatial dampening caused by coherence angle, defocus and spherical aberration (Tiemeijer et al., 2012). Additionally, the low energy spread helps reduce signal dampening caused by temporal coherence,

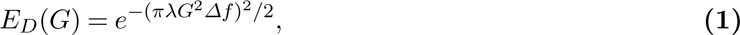

**Figure 1.**
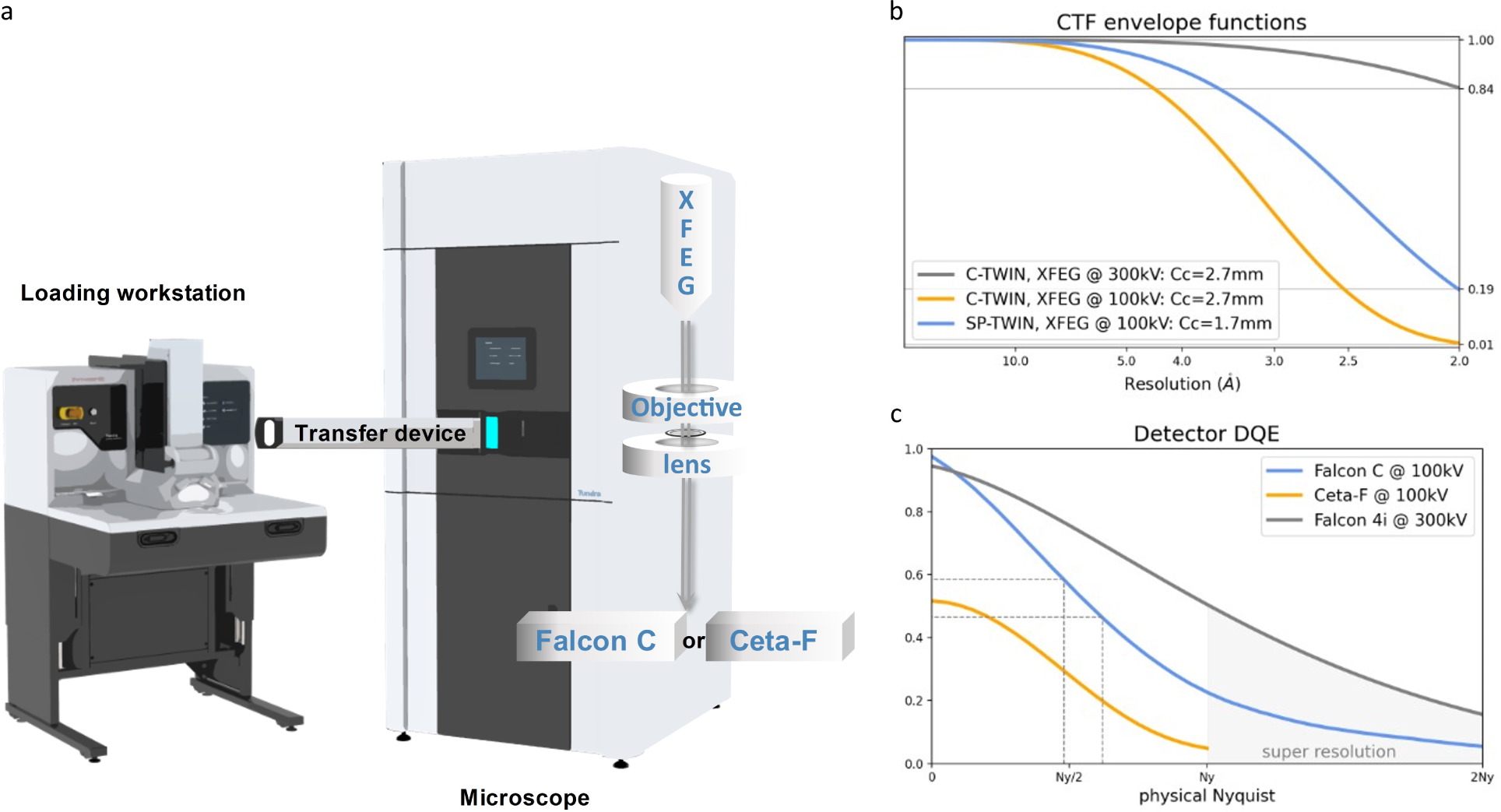
Cryo-EM at 100 keV. (**a**) Schematic overview of Tundra cryo-EM system equipped with a 100 kV XFEG, SP-TWIN objective lens and a choice of Ceta-F or Falcon C detector. The diagram also shows the loading workstation and cryo grid transfer device. (**b**) The envelope functions of different system configurations; XFEG at 100kV with C-TWIN (yellow) has a C_C_ of 2.7mm and C_S_ of 2.7 mm, XFEG at 100 kV with SP-TWIN (blue) has a C_C_ of 1.7 mm and C_S_ of 1.6 mm, and XFEG at 300 keV with C-TWIN lens. (**c**) Detective Quantum Efficiency (DQE) curves for Ceta-F (yellow) and Falcon C (blue) at 100 keV along with Falcon 4i (grey) at 300 keV.

where 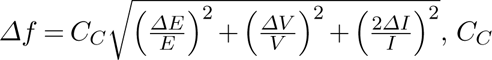 is the total chromatic aberration coefficient, 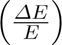 is the relative RMS energy spread of the electron gun, 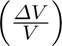 is the relative RMS instability of the high tension, and 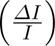 is the relative RMS instability of the objective lens current (Tiemeijer et al., 2012). Due to a longer wavelength and lower accelerating voltage at 100 keV (Supplementary Table S1), the signal still dampens significantly faster compared to higher accelerating voltages, for example, 300 keV (Figure 1b).

Furthermore, to improve the high-resolution signal, instead of using a regular objective lens with a pole piece gap of 11 mm and focal length of 3.5 mm (C-TWIN), the Tundra is equipped with an objective lens that has a pole piece gap of 7 mm and focal length of 2.3 mm (SP-TWIN). The SP-TWIN lens leaves enough space for the cryo box, an anti-cryo contamination device, which reduces water contamination on the sample and allows long data collection without decreasing sample and image quality. The SP-TWIN lens has an improved chromatic aberration coefficient, C_C_ of 1.7 mm, while C-TWIN has a C_C_ of 2.7 mm. Since C_C_ is a major contributor to signal improvement, with the smaller C_C_ of the SP-TWIN, the signal from 100 keV electrons at 2 Å resolution is improved by more than tenfold (from 1% to 19%, as shown in Figure 1b), although it remains substantially lower than the signal achieved at 300 keV.

In addition to these improvements to the electron source and electron optics, the Tundra microscope has been equipped with a new sample exchange mechanism, which we refer to as semi-automated loader. A detached loading workstation can house multiple sample grids in a fixed autoloader cassette, maintaining them under liquid nitrogen temperature until they are transferred to the microscope for imaging. Grid transfer to the microscope column involves a newly developed transfer device wherein all grid manipulations, including retrieval from the slot, placement into the transfer device, and subsequent transfer into the microscope, are performed automatically (Figure 1a). The only manual step is to detach the transfer device from the loading workstation and insert it into the microscope. During the transfer, samples are passively cooled by materials with high thermal mass; and the transfer device maintains vacuum conditions (10^-3^ Pa). The vacuum protects the cryo sample in cryo condition since the vacuum slows down conduction and convection providing several advantages over side-entry holders. First, the vacuum minimizes the accumulation of hexagonal ice contamination on the sample, and reduces the contamination of the microscope column during the sample transfer. Next, this system prevents vacuum crashes during sample insertion, which is a common occurrence with side-entry holders. After imaging, samples are also unloaded first under vacuum and then directly into liquid nitrogen, this also enables grid retrieval under cryogenic conditions with minimal ice contamination. Consequently, grids imaged on the Tundra can be further used for higher-resolution data collection using cryo-TEMs like the Krios and Glacios.

Detectors play a crucial role in the overall cryo-EM performance. While recent work demonstrated the feasibility of high-resolution cryo-EM structure determination at 100 keV (Naydenova et al., 2019; McMullan et al., 2023) no “large area” imaging detectors have been specifically designed for 100 keV electrons. To address this, we developed two new detectors, the Ceta-F CMOS detector with dose fractions and the Falcon C DED. The Ceta-F is a 4096 x 4096-pixel scintillator-based CMOS detector, which operates at a frame rate of 14 frames per second (fps). The frames are saved to the disk as individual dose fractions similar to a DED implementation, allowing for correction of beam-induced motion and dose weighting to compensate, in the post-processing pipeline, for the radiation damage. Applying a correlated double sampling (CDS) algorithm led to the dark noise reduction by a factor of two. Moreover, by simplifying the pixel design, each electron creates 4x counts when compared to the Ceta-16M, while the noise stays the same (Malínský et al., 2021). As a result, the detective quantum efficiency (DQE) at the low flux normally used for data collection (*∼*14e^-^/pixel/s), is improved to about 5% at Nyquist, around 26% at half Nyquist, and 52% at zero spatial frequency (Figure 1c).

The comparatively lower DQE of the Ceta-F detector relative to DEDs prompted us to develop a new DED optimized for use at 100 keV - the Falcon C, which has a sensor size of 6 cm x 6 cm and a 14 µm-thick epilayer (Kuijper et al., 2015; McMullan et al., 2016). The silicon used in Falcon C is back-thinned to 30 µm to reduce transverse scattering and enhance the signal-to-noise ratio. Reset noise is minimized by multi-frame correlated double sampling (mfCDS) (Kuijper et al., 2015; Janssen et al., 2014), a technology that has been in place since the third-generation Falcon detector. The remaining noise primarily arises from variations in the height and shape of the charge distribution produced by individual incident electrons or events (Supplementary Figure S1). To mitigate this Landau noise, we employed an electron counting algorithm that estimates the central position of each electron event and applies a constant signal at the estimated position. The imaging performance of the counting detector depends on both the efficiency of electron event detection and the accuracy of position estimation. The Falcon C sensor consists of 2048 x 2048 detector pixels (4 megapixels), each measuring 28 µm x 28 µm. The larger pixel size in combination of a thick epilayer enables a clear discrimination between noise and electron events. As shown in Supplementary Figure S1a, the dark noise follows a Gaussian distribution with a standard deviation of 0.36 DN (Camera internal digital unit), while electron events exhibit a Landau distribution with a peak at 7 DN.

The clear separation between signal and noise results in less than 0.23% of the electron events that fall below the noise level (false negatives) and less than 0.004% of the noise instances that are incorrectly identified as electron events (false positives). The average size of an electron event is represented by 2 x 2 pixels (Supplementary Figure S1b), enabling super-resolution imaging to be accomplished by estimating event positions using 2 x 2 subpixels, resulting in the generation of 4096 x 4096 pixels images after counting. This pixel design results in a high detective quantum efficiency (DQE) reaching 96% at zero spatial frequency, around 56% at half Nyquist frequency and approximately 22% at physical Nyquist frequency (Figure 1c). Additionally, the Falcon C operates at a frame rate of 250 frames per second (fps), yielding more than 85% detection efficiency at fluxes of up to 20 electrons/pixel/second (eps) (Supplementary Figure S1c).

To evaluate the performance of the Tundra microscope for biological structure determination, we collected data for four different protein samples using the Ceta-F detector and eight protein samples using the Falcon C detector. We benchmarked mouse apoferritin, Thermoplasma acidophilum 20S (T20S) proteasome, human *γ*-Aminobutyric acid type A (GABA_A_) receptor, and human haemoglobin on both detectors with the same microscope. One grid per sample was used when collecting data for comparative datasets. On the grid, areas of similar ice thickness were selected by an experienced microscopist. In addition to the reported resolution according to the Fourier Shell Correlation (FSC) at 0.143, we also estimated B-factors to assess the instrumentation and dataset quality (Rosenthal and Henderson, 2003). Unsurprisingly, apoferritin, which has a 24-fold symmetry and exceptional structural stability, was resolved at the highest resolution using both detectors. Two apoferritin datasets with either a C-TWIN or a SP-TWIN objective lens, paired with a Ceta-F detector were acquired. The SP-TWIN lens setup yielded a marked improvement in imaging quality resulting in a higher resolution, 2.6Å reconstruction with fewer particles with a B-factor of 174 Å^2^ compared to the C-TWIN dataset which required 1.5x the number of particles to achieve 3.0 Å resolution with a B-factor of 234 Å^2^ (Supplementary Table S2). Given the observed improvements associated with the SP-TWIN lens, all subsequent experiments were performed on the microscope with the SP-TWIN lens.

For Falcon C, we experimented data collection schemes at three different magnifications (0.55 Å/pix, 0.7 Å/pix and 0.9 Å/pix) to attempt to optimize the field of view, number of particles and resolution (Supplementary Figure S1). Initially, we processed data from the first 4 hours with pixel sizes of 0.7 Å/pix. From the 1,000 movies only, a reconstruction at 2.3 Å resolution from approximately 46,000 particles was obtained (Supplementary Table S3). When we expanded datasets to include around 5,000 movies per dataset (approximately 20 hrs of collection time) and processed it with RELION 4.0.1 including Bayesian polishing (Zivanov et al., 2022), we were able to achieve higher resolution reconstructions. Specifically, we obtained 2.2 Å resolution with 124,162 particles for the dataset with pixel size of 0.55 Å (Figure 2c, f, g) and 2.1 Å with 249,808 particles for the dataset with pixel size of 0.7 Å (Figure 2a, b, d, f, g). The dataset with 0.9 Å/pix also produced a 2.1 Å map, but it required 383,412 particles (Figure 2e, f, g, Supplementary Table S4). Once again, we assess the quality of the datasets collected at different magnifications by estimating their B-factors. Datasets from 0.55 Å/pix and 0.9 Å/pix resulted in the B-factors of 102.8 Å^2^ and 114.3 Å^2^ respectively (Figure 2h). Interestingly, the dataset of 0.7 Å/pix yielded slightly better results with B-factors of 95.7 Å^2^ (Figure 2h).

**Figure 2.**
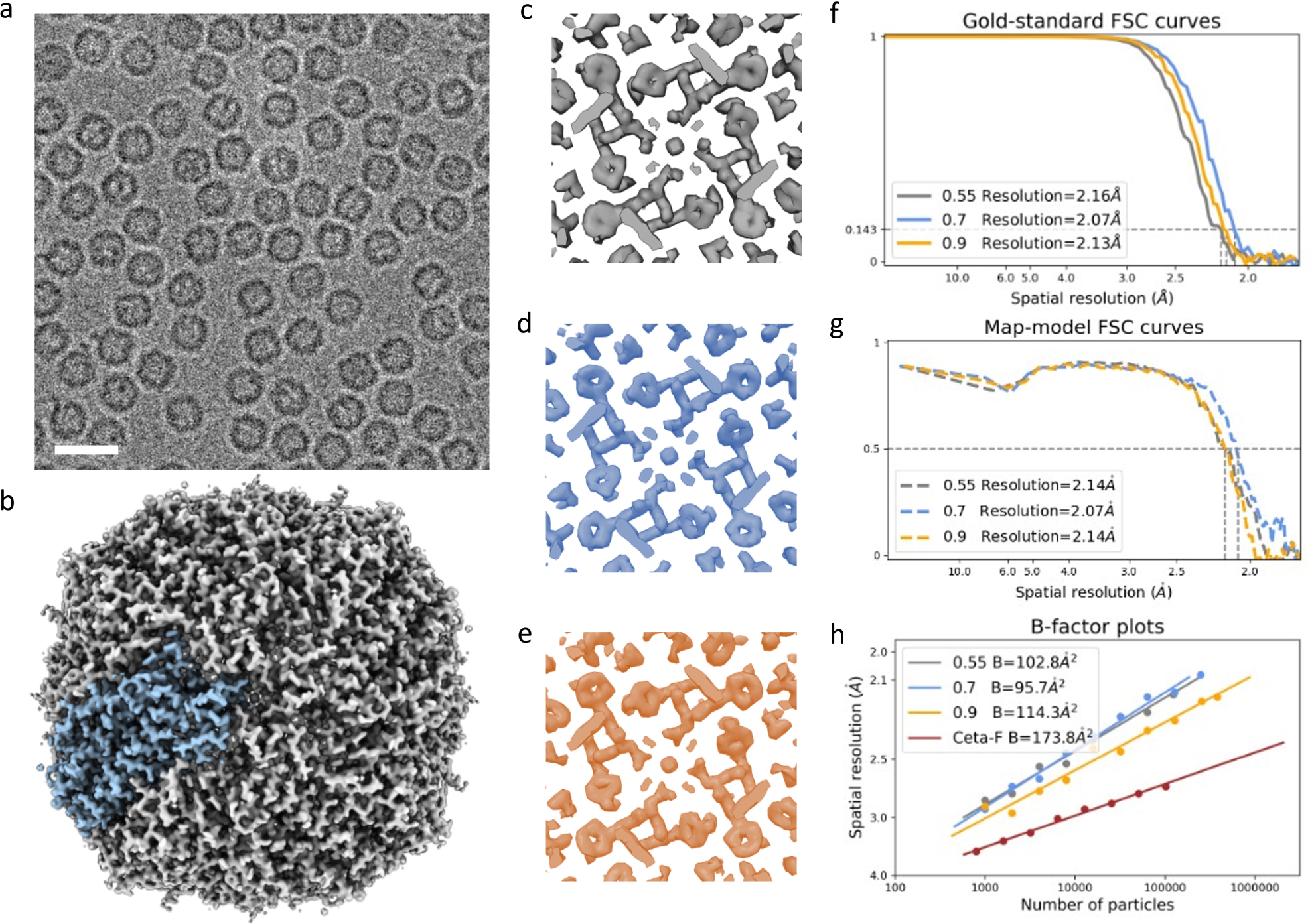
Apoferritin reconstructions imaged at 100 keV. (**a**) Representative motion-corrected and dose-weighted image of apoferritin sample in vitreous ice with 0.7 Å/pix (scale bar – 20 nm). (**b**) Three-dimensional reconstruction of apoferritin with octahedral symmetry, highlighting the asymmetric unit in blue. (**c-e**) Examples of high-quality density at 0.55, 0.7 and 0.9 Å/pix magnification respectively.(**f**) Gold-standard Fourier Shell Correlation (FSC) curves comparing the two independently refined half-maps of the datasets collected at three different magnifications: 0.55 Å/pix (grey), 0.7 Å/pix (blue), and 0.9 Å/pix (yellow). Resolutions are estimated using the FSC = 0.143 criterion. (**g**) Map-model Fourier Shell Correlation (FSC) curves comparing the experimental maps and the calculated model maps of the three datasets: 0.55 Å/pix (grey), 0.7 Å/pix (blue), and 0.9 Å/pix (yellow). Resolutions are estimated using the FSC = 0.5 criterion. (**h**) B-factor plots for four different datasets, including three collected at varying magnifications with the Falcon C detector (0.55 Å/pix in grey, 0.7 Å/pix in blue, and 0.9 Å/pix in yellow), and one dataset at a magnification of 0.75 Å/pix collected with Ceta-F (red).

The structure of the T20S proteasome, a 700 kDa protein complex with a D7 symmetry, was previously determined to a resolution of *∼*2.3 Å using a bottom-mounted Gatan K2 Summit DED on a 200 kV Talos Arctica (Herzik et al., 2017). A *∼*2.1 Å resolution structure was also reported at 200 keV using Glacios with the Selectris X energy filter and Falcon 4 detector (Koh et al., 2022). We determined the structure of T20S to *∼*3.0 Å resolution on Tundra with the Ceta-F detector using 183,486 particles (Supplementary Figure S1d-f). For the data collection on Falcon C, we followed a similar approach as described for apoferritin, although we only collected datasets at two different magnifications. Data processing the first four hours of data at 0.7 Å/pix magnification yielded a *∼*3.1 Å resolution map with D7 symmetry applied. We were able to further improve the resolution by increasing the size of the dataset. With a similar number of particles as in the Ceta-F dataset, we obtained a *∼*2.7 Å resolution map using Falcon C (Figure 3a-b, f). The lower magnification dataset (0.9 Å/pix), which yielded more particles due to the lower magnification and larger field of view, nonetheless yielded a lower resolution reconstruction of *∼*2.9 Å (Supplementary Table S4) likely due to the reduced DQE of the detector at these settings. For example, 2.9 Å is equivalent to 0.48 Nyquist when using a pixel size of 0.7 Å/pix, while the same resolution, 2.9 Å, is equivalent to 0.62 Nyquist when using a pixel size of 0.9 Å/pix. The DQE at 0.48 Nyquist is 14% higher than at 0.62 Nyquist on Falcon C, 59% vs. 46% (Figure 1c).

**Figure 3.**
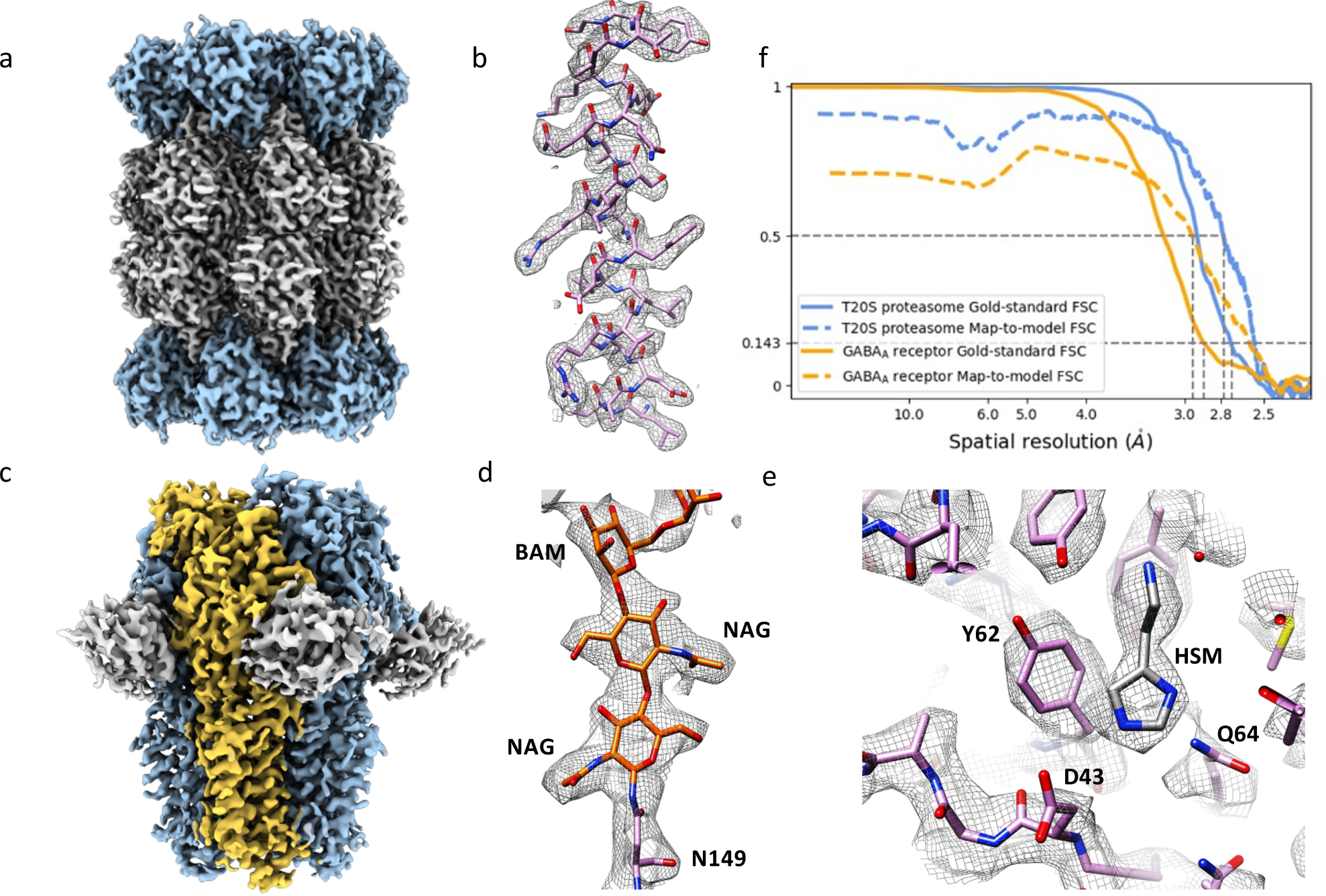
Cryo-EM reconstructions of the 20S proteasome and GABA_A_ receptor from Falcon C detector. (**a**) A 2.7 Å cryo-EM map of 20S proteasome with D7 symmetry. (**b**) Example of an alpha-helix, represented as sticks, with corresponding EM density (grey mesh, zoned 2 Å within atoms) showing clearly resolved side-chain densities. (**c**) A 2.8 Å cryo-EM map of GABA_A_ receptor with C5 symmetry, viewed parallel to the plasma membrane space (side view). The density of one subunit of the homopentamer is highlighted in yellow, and the bound nanobody domain Mb5 is shown in grey. (**d**) Visualization of the N-acetyl glucosamine (NAG) moieties attached to Asn149. (**e**) An overview of map quality at the agonist binding pocket, depicting bound histamine (HSM). (**f**) Gold-standard FSC curve and map-model FSC curve corresponding to the map in (a) and (c).

To further evaluate the resolving capabilities of 100 keV on more challenging samples, we focused on a *∼*200 kDa human membrane protein, the *β*3 homo-pentameric GABA_A_ receptor bound to the Mb25 megabody and its small molecule agonist, histamine (Miller and Aricescu, 2014). GABA_A_ receptors are a large family of pentameric ligand-gated chloride channels that carry the primary target sites for a wide range of clinically relevant drugs including general anaesthetics, benzodiazepines, barbiturates, and neuroactive steroids (Sieghart and Savić, 2018). With the Ceta-F, we were able to determine the GABA_A_ structure to *∼*3.4 Å resolution from 92,997 particles, with a C5 symmetry imposed (Supplementary Figure S1g-i). Using the same magnification and five-fold fewer particles (17,526 particles) from 1,000 movies collected in four hours, we determined the structure of this receptor to *∼*3.3 Å using the Falcon C detector (Supplementary Table S3). Furthermore, by using the particles from the full dataset collected on Falcon C, we improved the resolution to *∼*2.8 Å with a similar number of particles as were used for the Ceta-F dataset (Figure 3c, f; Supplementary Table S5). At this resolution, we were able to resolve the individual glycan units attached to on Asn149 and the histamine molecules bound within the five orthosteric pocket (Figure 3d-e).

We also show that the Tundra enables high-resolution reconstruction of specimens that are at the lower size limit of resolvability using current state-of-the-art cryo-EM instrumentation. Imaging haemoglobin, a 64 kDa soluble protein complex with C2 symmetry, with the Ceta-F, produced 2D class averages containing secondary structure features (Figure 4a). Subsequent iterative 3D classifications yielded a particle set that led to a *∼*8.1 Å resolution reconstruction, which was of sufficient quality to clearly discern the molecular envelope and identify the positions of alpha helices (Figure 4b, d). At this resolution, it is possible to detect large conformational changes, such as in this case, differentiating between conformers of oxyhaemoglobin and deoxyhaemoglobin (Naydenova et al., 2019). Notably, when we collected data on the same sample using the Falcon C detector, the 2D class averages contained more detailed secondary structure features (Figure 4e), an improvement we attribute to the enhanced signal-to-noise ratio of Falcon C. The 3D resolution of the reconstruction was also significantly improved over the Ceta-F dataset, as we were able to determine the structure to *∼*5.0 Å resolution from 66,157 particles with C2 symmetry imposed. In this reconstruction, the alpha helices were clearly distinguishable, and the protein backbone could be confidently traced. Moreover, clear density can be observed for the heme groups (Figure 4f,h).

**Figure 4.**
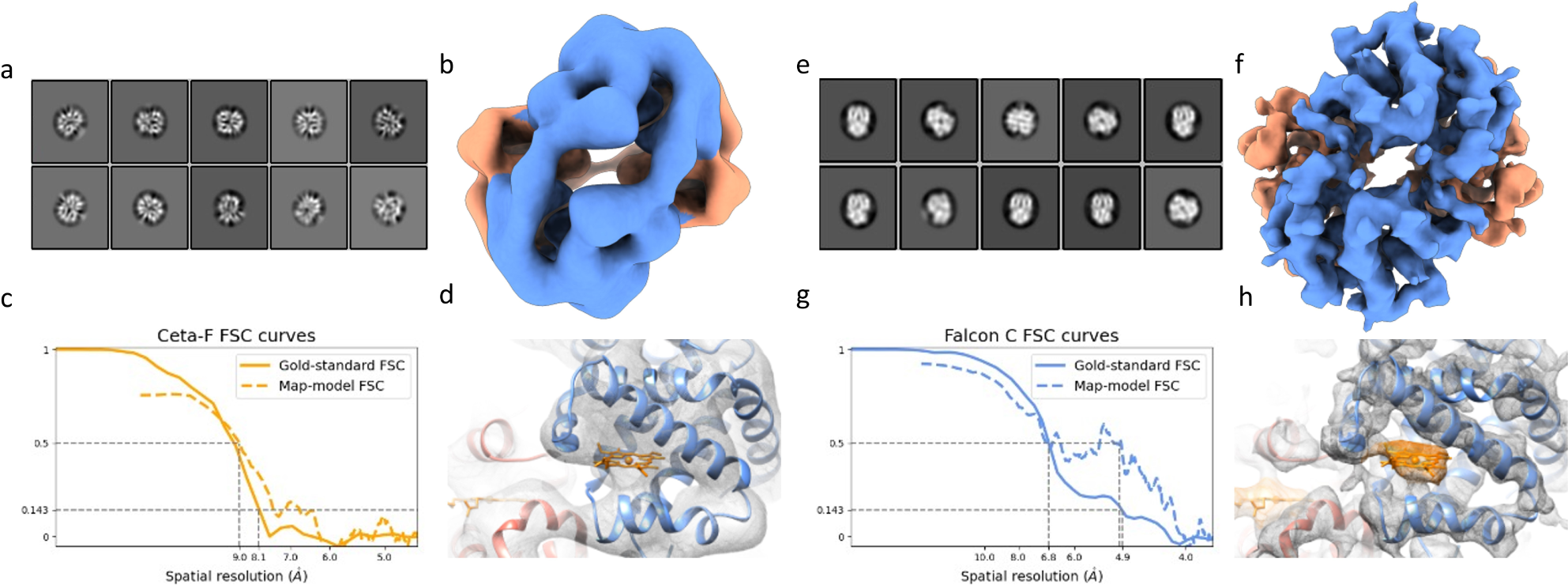
Structure of haemoglobin determined at 100 keV using Ceta-F and Falcon C detectors. (**a**) Reference-free 2D class averages of haemoglobin from the data collected on the Ceta-F detector. (**b**) An 8.1 Å reconstruction of haemoglobin obtained with the Ceta-F detector. (**c**) Gold-standard FSC curve and map-model FSC curve corresponding to the map in (b). (**d**) Example of EM density with a fitted model showing clear densities for alpha-helices. (**e**) Reference-free 2D class averages of haemoglobin from the data collected on the Falcon C detector. The class averages from Falcon C have a higher signal-to-noise ratio, resulting in sharper features for secondary structure elements compared to Ceta-F data in (a). (**f**) A 5.0 Å reconstruction of haemoglobin obtained with the Falcon C detector. (**g**) Gold-standard FSC curve and map-model FSC curve corresponding to the map in (f). (**h**) Example of EM density with a fitted model showing clear densities for alpha-helices and the bound heme group.

Encouraged by the above results, we further investigated the performance of the Tundra microscope with the Falcon C detector with other small (for cryo-EM standard) protein. Our aim was to test whether the Tundra with this direct electron detector, specifically optimized for 100 keV, serves not only as a screening microscope but also as an effective tool for determining high-resolution single particle cryo-EM structures. The first of these samples, *∼*150 kDa rabbit muscle aldolase, was previously resolved to *∼*2.1 Å resolution using a Talos Arctica microscope operating at 200 keV (Wu et al., 2020). Remarkably, with just 1,000 movies and four hours of data acquisition, we obtained a *∼*3.4 Å reconstruction from 14,675 particles (Supplementary Table S3). By incorporating more data and iterative 3D classification and refinement, with D2 symmetry, followed by Bayesian particle polishing in RELION 4.0.1 (Zivanov et al., 2022), we further improved the resolution to *∼*2.8 Å using 143,939 particles (Figure 5a-c). The final reconstruction exhibits well-resolved backbone density and clearly defined side chain densities across most of the molecule. Next, we looked at human CDK-activating kinase (CAK), a *∼*85 kDa heterotrimeric protein complex formed by CDK7, cyclin H, and MAT1, which acts as a master regulator of cell growth and division (Cushing et al., 2024). Previously, we reported the structure of the CDK-cyclin-module of the human CAK in its nucleotide-bound form at *∼*1.8 Å and *∼*2.3 Å resolution using 300 kV and 200 kV microscopes equipped with Selectris energy filter and Falcon 4i direct electron detector (Cushing et al., 2024). Data collected using the Falcon C on the Tundra enabled us to resolve the structure of CAK to *∼*4.0 Å from 219,717 particles (Supplementary Table S5), which was of sufficient resolution to be able to trace the backbone of the complex. Furthermore, we were able to distinguish the density for the bound AMPPNP and some of the larger side chains (Figure 5d-f).

**Figure 5.**
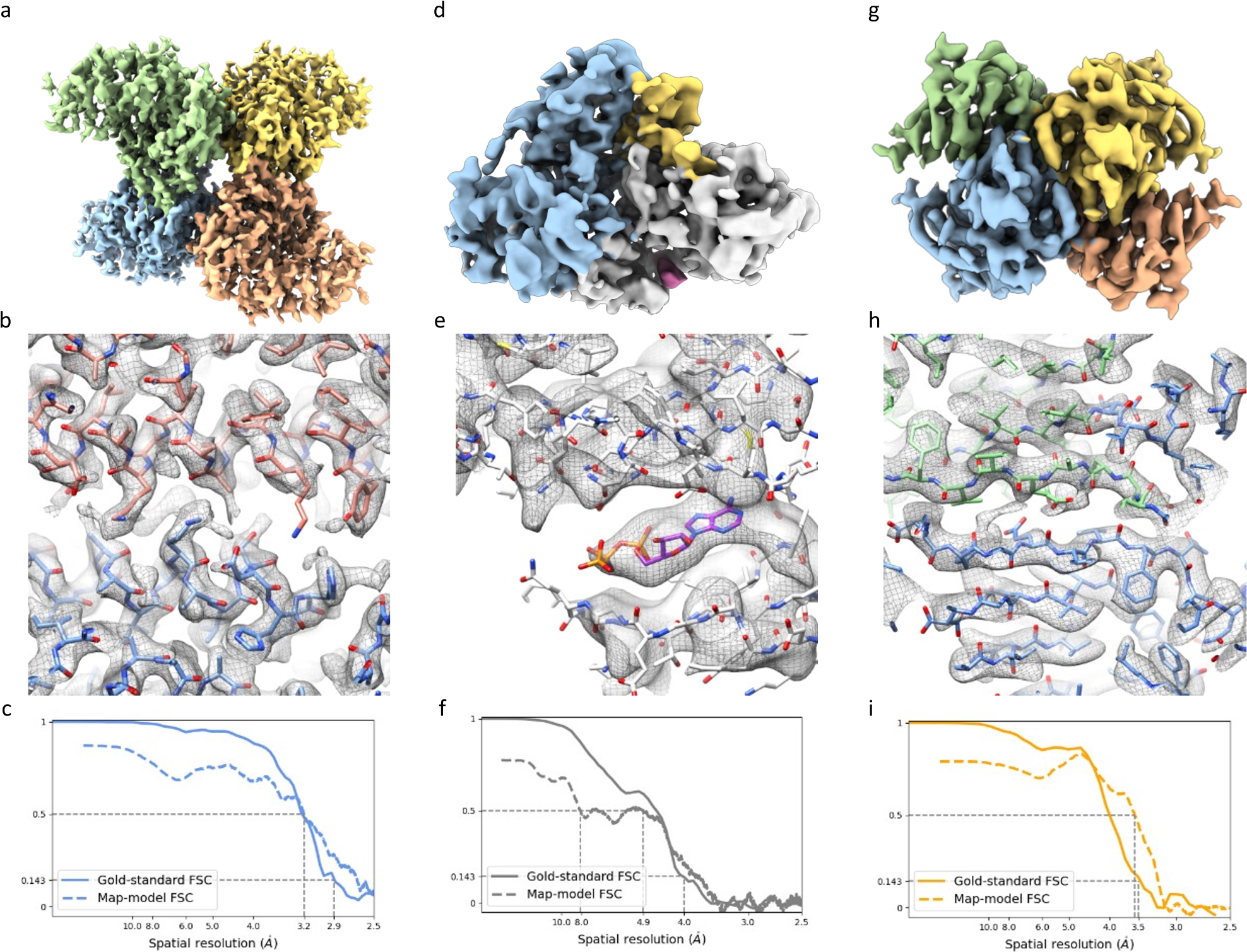
Cryo-EM structures of aldolase, CAK and transthyretin determined at 100 keV on Falcon C detector. (**a**) cryo-EM reconstruction of rabbit muscle aldolase at 2.8 Å resolution coloured by subunits with D2 symmetry. (**b**) A zoomed-in region of the EM density (grey mesh) displaying alpha-helices, fitted with an atomic model in stick representation.(**c**) Gold-standard FSC curve and map-model FSC curve corresponding to the aldolase map in (a). (**d**) A 4.0 Å cryo-EM map of CAK with distinctly coloured CAK subunits (cyclin H blue, MAT1 yellow, CDK7 grey, AMP-PNP purple). (**e**) A close-up view of the density for bound AMP-PNP in the CDK7 active site pocket. (**f**) Gold-standard FSC curve and map-model FSC curve corresponding to the CAK map in (d). (**g**) A 3.4 Å reconstruction of transthyretin coloured by subunits with D2 symmetry. (**h**) A close-up view of the EM density in the region of beta-strands, showing clear strand separation and fitted side chains. (**i**) Gold-standard FSC curve and map-model FSC curve corresponding to the transthyretin map in (g).

We next looked at a smaller protein assembly, the human transthyretin (TTR) homotetramer, which comprises four identical polypeptide chains with D2 symmetry, collectively amassing 55 kDa. TTR is a conserved transport protein that is present in vertebrates and is produced in the liver, choroid plexus, and retinal pigment epithelium (Sanguinetti et al., 2022; Liz et al., 2020) and facilitates the transport of the thyroid hormone thyroxine. It also plays a role in carrying vitamin A by forming a complex, with retinol binding protein (RBP) (Liz et al., 2020). Cryo-EM structures of this tetrameric form of TTR were solved to 2.7 Å using a 200 kV Talos Arctica, revealing inherent asymmetry in the tetrameric architecture and previously unobserved conformational states (Basanta et al., 2024).

We successfully determined the structure of TTR at a resolution of *∼*3.5 Å from 131,895 particles with D2 symmetry imposed. Each monomer of TTR predominantly consists of beta-strand structures with a single, short alpha-helical segment. At this resolution, we could clearly distinguish individual beta-strands, and visualize most side chains with the map (Figure 5g-i). To our knowledge, this is the highest resolution structure that has been achieved by single particle analysis of such a small protein using a 100 kV TEM.

Finally, at the other end of the size spectrum, we also explored a larger specimen: an Adeno-associated virus (AAV9) capsid. This virus like particle (VLP) weighs 3.9 MDa and possesses icosahedral symmetry. We collected data at 0.7 Å/pix magnification, which corresponds to a limited field view for such large particles on a 2K x 2K detector. Nevertheless, we could still capture on average 13 VLPs per movie. Iterative 2D and 3D classification yielded a subset of 20,369 particles from 6,689 movies that, after 3D refinement with icosahedral symmetry followed by Bayesian polishing, resulted in a *∼*2.8 Å reconstruction of its structure (Supplementary Figure S1).

## Discussion

Our results demonstrate the capabilities and potential of a 100 kV electron microscope equipped with a field-emission gun (FEG), a low C_C_ objective lens and two different types of detectors - the Ceta-F, a scintillator-based CMOS detector and the Falcon C, a direct electron detector - for high-resolution imaging and reconstruction of vitrified biological samples. To truly make cryo-EM accessible to all researchers, and in particular, to engage new users of the method, more affordable microscopes and automation protocols for sample handling, data collection and data processing are highly desirable. The semi-automated loader offers a new and unique way for frozen hydrated sample exchange. Compared to traditional side-entry systems, this technology greatly enhances the convenience and efficiency of sample screening, which is a critical aspect of any cryo-EM project. The semi-automated loader allows data collection for a long duration not only without the need to manually refill liquid nitrogen but also without the holder drift introduced by liquid nitrogen. Further, the semi-automated loader also allows sample retrieval and reuse after the long-duration data collection. The Tundra microscope is a fully integrated computer-controlled system with the capacity to perform automated data collection on biological samples. With the aberration-free image shift method (Cheng et al., 2018; Zivanov et al., 2018) integrated in the EPU software, we were able to collect data with a throughput of 200-300 movies per hour. Data processing workflows are now widely accessible through the continued efforts of developers of software packages such as RELION (Zivanov et al., 2022) and CryoSPARC (Punjani et al., 2017), enabling on-the-fly processing and high-resolution reconstruction of protein samples with minimal user intervention.

Our results demonstrate the ability of this affordable 100 keV cryo-TEM to obtain high-resolution cryo-EM structures when paired with either of two detectors. We achieved a remarkable *∼*2.6 Å resolution for apoferritin using a scintillator-based CMOS detector, and the implementation of a new direct electron detector improved our resolution to *∼*2.1 Å. We also extended our investigation to encompass a variety of challenging biological samples, including a human membrane protein, the GABA_A_ receptor, haemoglobin, rabbit muscle aldolase CAK, TTR, and AAV capsids. These diverse samples cover a range of molecular weights and symmetries, from small asymmetric proteins to large viral capsids. It is worth noting that our ability to accurately perform particle alignment of small samples such as haemoglobin is worse for datasets collected using the Ceta-F detector compared to the Falcon C detector, resulting in noisier class averages and lower resolution structures. This outcome is expected, considering that the Ceta-F detector relies on scintillator technology and consequently has a lower DQE. The lack of signal in the low to medium resolution limits the accuracy in matching the particles to the reference and thus results in low-resolution reconstructions. Ceta-F is nonetheless able to show molecular envelopes and thus would work well as a screening detector.

Despite the promising results obtained with our 100 kV electron microscope and direct electron detector, we acknowledge certain technical limitations. Although 100 keV has the benefit of more signal per unit of radiation damage when compared to 300 keV, this advantage diminishes quickly when the ice/sample thickness increases. Cryo-EM sample preparation is still fraught with numerous challenges that must be overcome to produce grids that preserve biological specimens in a thin layer of vitrified ice with random orientations. A majority of samples that are being imaged currently have significantly thicker ice than 30 nm, as many samples, especially membrane proteins, are better preserved in ice that is thicker than 30 nm (Peet et al., 2019). For such samples, 200 or 300 kV microscopes are more efficient.

Furthermore, the performance of the electron optics at 100 keV remain worse than those on 300 kV Krios instruments for example. The electron wavelength of at 100 keV is twice as long as at 300 keV (Supplementary Table S1). All electron optical aberrations, including C_C_, Ewald sphere, coma, and other higher order aberrations, have a larger impact at 100 keV than 300 keV (Tiemeijer et al., 2012). The effects from C_C_ and energy spread are shown in Figure 1b. The effects of Ewald sphere, coma and other higher-order aberrations are theoretically correctable using single particle reconstruction packages such as RELION (Zivanov et al., 2018, 2022). However, we lack a clear understanding of the residual aberrations that persist after correction and their effects on resolution at 100 keV. Further investigation is needed to determine the resolution limit from high-order aberrations at 100 keV. The signal generated from the elastically scattered electrons at 100 keV, although more efficient per radiation damage for thin ice/sample, suffers from significant loss due to the current limitations of electron optics (McMullan et al., 2023).

Additionally, due to their limited imaging power, 100 kV microscopes are not suited for thick specimens. Cell or tissue lamellae of 50-200 nm are inherently too thick, and the stage tilting required for tomographic data collection further exacerbates this problem.

Finally, detectors at 100 keV are still not as optimal as those currently available for 300 keV. The low speed of electrons accelerated at 100 keV requires a larger physical pixel size at the detector, which limits the number of pixels that can be positioned on a detector of a given dimension. For example, the Falcon C is a 2048 x 2048-pixel detector because of its larger pixel size, while the Falcon 4i is a 4096 x 4096-pixel detector. An image from the Falcon 4i has a field of view four times as large as an image from the Falcon C when using the same pixel size on the object scale, resulting in decreased throughput of the Falcon C relative to the Falcon 4i. While the issue of throughput could be overcome by generating a 4k x 4k detector with this larger detector pixel size, such a detector would be physically much larger and prohibitively expensive, which is against the concept of a lower-cost and easier-to-access cryo-TEM. To achieve the full potential of cryo-EM at 100 keV, counting algorithms should be further improved. A possible avenue could be neural network algorithms trained for the dedicated sensor design operating at 100 keV.

We conclude that while 300 kV TEMs deliver the highest throughput and the highest attainable resolution images for cryo-EM reconstructions, the 100 kV TEMs can be used for fast sample screening and can additionally provide high-quality datasets and high-resolution cryo-EM reconstruction, especially for symmetric samples. Our work paves the way for further advancement of cryo-EM instrumentation and high-resolution investigation of macromolecules.

## Methods

### Protein purification and cryo-EM grid preparation

### Apoferritin

A frozen aliquot of 7 mg/ml mouse apoferritin in 20 mM HEPES pH 7.5, 150 mM NaCl, 1 mM dithiothreitol (DTT) and 5% trehalose, which we received from the Kikkawa laboratory at Tokyo University, was thawed at room temperature and cleared by centrifugation at 10,000 g for 10 min. The supernatant was diluted to 5 mg/ml with 20 mM HEPES pH 7.5 150 mM NaCl, and 3 µl of the diluted sample was applied onto glow-discharged R1.2/1.3 300 mesh UltrAuFoil gold grids (Quantifoil) (Russo and Passmore, 2016) for 30 s and then blotted for 5 s before plunge-freezing the grids into liquid ethane cooled by liquid nitrogen. Plunge-freezing was performed using a Vitrobot Mark IV (Thermo Fisher Scientific) at 100% humidity and 4 °C.

### GABA_A_ receptor production, purification and nanodisc reconstitution

The homopentameric GABA_A_ receptor *β*3 was expressed and purified as previously described (Miller and Aricescu, 2014; Nakane et al., 2020; Uchański et al., 2021). Briefly, a construct encoding the human GABA_A_ receptor *β*3 subunit (Uniprot ID P28472) with the M3-M4 loop replaced by the SQPARAA sequence and an N-terminus SBP tag, was cloned into the pHL vector (Aricescu et al., 2006) and transiently transfected into Expi293F cells. Cell pellets from 1L culture (12 ml) were resuspended in 20 ml PBS pH = 7.4 supplemented with one cOmplete Protease Inhibitor Cocktail EDTA-free tablet (Roche). The suspension was supplemented with 3.5 ml of 10% (w/v) lauryl maltose neopentyl glycol (LMNG) stock in water and incubated for 1 hour at 4 °C with gentle rotation to solubilize cellular membranes. The insoluble material was removed by centrifugation at 12,500 g, 4 °C, for 15 minutes, and the supernatant was incubated for 2 hours with 300 µl Streptavidin beads (Thermo Fisher Scientific) at 4 °C. The beads were washed in batch, three times, with 50 mL PBS pH = 7.4, 0.1% (w/v) LMNG to remove impurities. The receptor was then reconstituted on beads in MSP2N2 nanodiscs (Addgene ID: 29520) in the presence of a 85:15 POPC:bovine brain extract (Sigma-Aldrich; Type I, Folch Fraction I; prepared as a 20 mg/ml stock in 3% DDM) mix, as described previously (Masiulis et al., 2019; Laverty et al., 2019). Reconstituted receptors were eluted with 200 microliters 2.5 mM biotin in PBS pH = 7.4 and concentrated to 1.2 mg/mL using an Amicon® Ultra Centrifugal Filter (100 kDa MWCO). Prior to grids freezing, 1 mM histamine and 3 mM Mb25 (Uchański et al., 2021) were added to the sample which was incubated for 1h on ice. Afterwards, 3.5 µL sample was applied to a freshly glow-discharged (PELCO easiGlow, 30 mA for 120s) 0.6/1 UltrAuFoil grid (Quantifoil) (Russo and Passmore, 2016), which was blotted for 3 s and vitrified in liquid ethane cooled by liquid nitrogen by using a Vitrobot Mark IV (Thermo Fisher Scientific) plunger at 100% humidity and 4 °C.

### T20S Proteasome

Thermoplasma acidophilum 20S proteasomes were recombinantly expressed in Escherichia coli and purified as described in (Zwickl et al., 1992). 4.5 µl of the purified T. acidophilum 20S proteasome sample was applied onto a glow-discharged 200 mesh Quantifoil R2/1 or R1.2/1.3 copper grid, and then blotted for 4.5 s before plunge-freezing with liquid ethane cooled by liquid nitrogen. Samples were vitrified using a Vitrobot Mark IV (Thermo Fisher Scientific) set to 4 °C and 100% humidity.

### Haemoglobin

Lyophilized Human haemoglobin (Sigma-Aldrich) was resuspended in 50 mM HEPES pH 7.4 to a final concentration of 3mg/ml and centrifuged at 10,000 g for 10 min at 4°C. 3ul sample was applied onto glow-discharged R1.2/1.3 300 mesh UltrAuFoil gold grids (Quantifoil) (Russo and Passmore, 2016), blotted for 5s and plunge frozen into liquid ethane using Vitrobot Mark IV (Thermo Fisher Scientific) at 100% humidity and 4 °C.

### Adeno-associated Virus 9

HEK293F cells were transiently transfected with a mixture of three plasmids: the transfer plasmid DNA containing GFP gene, Rep/Cap6 plasmid DNA and helper plasmid DNA. All three plasmids were mixed at a mass ratio of 1:3:4 and transferred to a vial containing CTA Viral-Plex complexation buffer at 4-8 °C. The CTS AAV-MAX transfection reagent was added to the CTS AAV-MAX transfection booster, incubated for 10 minutes and transferred to the mixture containing the plasmid DNA and Viral-Plex complexation buffer. The reaction mixture was further incubated for 20 minutes without agitation and added to the cell culture. 72 hours post-transfection, the cells were harvested by addition of 10x AAV-MAX lysis buffer together with 2 mM MgCl2 and 90 U/mL of Pierce Universal Nuclease to digest any non-encapsulated nucleic acid. The cell lysate was clarified by filtration. The clarified cell lysate was concentrated 20x using hollow-fiber tangential flow filtration (HF-TFF). The material was buffer exchanged with affinity equilibration buffer (100 mM Tris, 500 mM NaCl, 0.01% Pluronic F-68 surfactant, pH 7.2) and then loaded onto a column packed with POROS CaptureSelect AAVX resin (Thermo Fisher Scientific) at a flow rate of 300 cm/hr. The unbound proteins were washed off the column with 5 column volumes of 3 different wash buffers (Wash 1: 50 mM Tris, 1.5 M NaCl, 0.01% Pluronic F-68 surfactant, pH 7.2, Wash 2: 50 mM Tris, 1.5 M urea, 0.01% Pluronic F-68 surfactant, pH 9.0, Wash 3: 50 mM Tris, 1% Tween^TM^ 20 surfactant, 0.01% Pluronic F-68 surfactant, pH 9.0). AAV6-GFP capsids were eluted using elution buffer (75 mM glycine, 50 mM NaCl, 500 mM arginine, 0.01% Pluronic F-68 surfactant, pH 2.5). The viral genome titers were determined using QX200^TM^ Droplet Digital PCR System (Bio-Rad). The purified AAV6 sample was concentrated to *∼*1.5-2 mg/ml by centrifugation at 10,000 g, 4 °C using 100 kDa MWCO PES concentrator (Amicon). 3 ul of concentrated sample was applied onto glow discharged R1.2/1.3 300 mesh holey carbon grids (Quantifoil), blotted for 6 s and plunge frozen into liquid ethane using a Vitrobot Mark IV (Thermo Fisher Scientific) at 100% humidity and 4 °C.

### Aldolase

Lyophilized aldolase from rabbit muscle (Sigma-Aldrich) was resuspended in 20 mM HEPES pH 7.4, 50 mM NaCl to a final concentration of 3mg/ml and centrifuged at 10,000g for 10 min at 4°C. 3 ul sample was applied onto glow-discharged R1.2/1.3 300 mesh UltrAuFoil gold grids (Quantifoil) (Russo and Passmore, 2016), blotted for 5 s and plunge frozen into liquid ethane using Vitrobot Mark IV (Thermo Fisher Scientific) at 100% humidity and 4 °C.

### Transthyretin

BL21 (DE3) competent E. coli (New England BioLabs, Ref *#*C2527H) were transformed with a pMMHa vector encoding the transthyretin gene. Transformed cells were selected by growing them on agar plates E. coli starter cultures were started by selecting a single colony of transformed cells from an agar plate containing 100 µg/mL ampicillin and grown at 37 °C. The colony was cultured in 50 mL LB media in a 250 mL shake flask containing 100 µg/mL ampicillin at 37 °C. The cultures were shaken at 37 °C until bacterial growth until the media appeared turbid. Expression cultures were prepared by inoculating a 1 L shake flask containing 100 µg/mL ampicillin with a 1:20 dilution of the starter culture, and grown at 37 °C until OD600 was approximately 0.5. Cultures were then induced with 1mM IPTG and incubated overnight at 30 °C while shaking. Cultures were harvested by centrifugation at 6,000 rpm for 30 minutes at 4 °C and the supernatants were removed. Pellets were resuspended with 50 mL Tris-Buffered Saline (TBS) with an added protease inhibitor tablet (Thermo Scientific Pierce Protease Inhibitor Tablets EDTA-Free, Ref *#*A32965), followed by probe sonication (3 times with Qsonica Q125; 3 minutes sonication / 3 minutes rest at 4 °C). Sample was centrifuged at 15,000 rpm for 30 minutes, the supernatant collected, and subjected to 50% ammonium sulfate precipitation (w/v). The sample was then stirred for 45 minutes at room temperature, and then the supernatant was collected after centrifugation at 15,000 rpm for 30 minutes at 4 °C. The supernatant was precipitated again with 90% ammonium sulfate (w/v) and stirred for 45 minutes at room temperature, and then centrifuged. The pellets were collected and dialyzed with 3,500 MWCO dialysis tubing (SnakeSkin Dialysis Tubing, Ref *#*68035) overnight against 25 mM Tris pH 8.0 buffer in a 4 °C cold room. The resulting protein mixtures were filtered using a low protein binding filter (Millipore Sigma), and purified through a Source 15Q anion exchange column (Cytiva) at a flow rate of 2 mL/min. Protein mixtures were injected at 5% Buffer A (25 mM Tris pH 8.0) and eluted using a 60-minute gradient up to 100% Buffer B (25 mM Tris/1.0 M NaCl) at room temperature. Eluates were subjected to further purification steps using a Superdex 75 gel filtration column. Transthyretin proteins were eluted with 10 mM sodium phosphate pH 7.6 / 100 mM KCl. The stabilizing A2 ligand was covalently bound in each of the TTR binding pockets, by setting up a reaction where 1 µL of a 1.5 mM solution of A2 in DMSO was added to a 19 µL of 1.7 mg/mL TTR stock solution. The A2 small molecule and TTR were incubated for 4 hours in the dark at room temperature. Due to the preferred orientation of TTR tetramers relative to the air-water interface when prepared on traditional holey carbon grids, TTR sample was frozen on prepared graphene grids, which were prepared using a previously described protocol [REF PMID: 37747197]. A single monolayer of graphene was applied to R1.2/1.3 UltrAuFoil Holey Gold grids (Russo and Passmore, 2016). The monolayer graphene was oxidized by exposure to UV/ozone for four minutes using the UVOCS T10x10 system. Cryo-EM samples were prepared using a manual plunger in a cold room that was maintained at 6 °C and 90% humidity. Grids were mounted on the plunger and 3 µL of sample was applied onto the graphene side of the grid and immediately blotted for 6 seconds (counting started after the blotted liquid spot on the filter paper stopped spreading) using a 1 x 6 cm piece of Whatman *#*1 filter paper. The paper was held parallel to the grid to establish full contact, and time was kept consistent using a metronome. The grid was plunged in liquid ethane at the same time the blotting paper was pulled back.

### Cryo-EM data collection

All cryo-EM data collection was performed on Tundra cryo-TEM (Thermo Fisher Scientific) operating at 100 keV using either Ceta-F detector or Falcon C detector. All datasets on Ceta-F were acquired in integration mode operating at 14 frames per second in CDS mode with the exception of apoferritin (C-TWIN) and haemoglobin that had Ceta-F operating at 18 frames per second in non-CDS mode, using EPU software (Thermo Fisher Scientific). For apoferritin (SP-TWIN), GABA_A_ receptor, and T20S proteasome data sets, a nominal magnification of 180,000x resulting in pixel size of 0.75 Å/pixel, total dose of 30-35 e-/Å^2^ with electron flux of 12.95-14.66 e^-^/pixel/s and exposure time of 1.1-1.3 s were used. Apoferritin (C-TWIN) data set was collected using 190,000x nominal magnification with pixel size of 0.73 Å, total dose of 40 e^-^/Å^2^, electron flux of 22 e^-^/Å and 1 second exposure time. For haemoglobin, -48 e^-^/Å^2^ with electron flux of 31 e^-^/pixel/s and exposure time of 0.51 s was used. All the Ceta-F datasets are in MRC format while Falcon C datasets are collected in EER format (Guo et al., 2020).

Falcon C datasets of mouse apoferritin, T20S proteasome and human GABA_A_ receptor were collected with a total dose of 40 e^-^/Å^2^. Rabbit muscle aldolase, CAK complex, haemoglobin and human transthyretin were imaged with a total dose of 50 e^-^/Å^2^. Nominal magnifications of 290,000x, 370,000x and 470,000x corresponding to 0.9 Å/pixel, 0.73 Å/pixel and 0.55 Å/pixel were used for apoferritin. Datasets for T20S proteosome and human GABA_A_ receptor were collected at 290,000x, 370,000x corresponding to 0.9 Å/pixel and 0.73 Å/pixel. Datasets for rabbit muscle aldolase were collected at 370,000x and 470,000x corresponding to 0.73 Å/pixel and 0.55 Å/pixel. The dataset for CAK complex and haemoglobin was collected at 290,000x (0.9 Å/pixel). The dataset for human transthyretin was collected at 470,000x (0.55 Å/pixel). A defocus range of 0.2 µm to 0.6 µm in steps of 0.2 µm was used for mouse apoferritin and T20S proteasome. All other datasets used a defocus range of -0.3 µm to -0.9 µm. Dose rates at parallel illumination conditions ranged from 6-8 eps resulting in exposure times of 1.7 to 4.9 seconds.

### Image processing

Gain-corrected Ceta-F MRC or Falcon C EER and gain files were first imported into RELION (Zivanov et al., 2022). Motion correcting the beam-induced motion in the movies was done via RELION’s implementation of the MotionCor2 algorithm (Zheng et al., 2017). The contrast transfer function was estimated with CTFFIND 4 (Rohou and Grigorieff, 2015) via the sums of power spectra obtained from fractions with an accumulated dose of 4 e^-^/Å^2^. Images with poor quality were filtered out based on the estimated resolution and figure of merit parameters from CTFFIND 4.

Particles from Ceta-F datasets were picked using RELION’s reference-based algorithm and subjected to several rounds of 2D classification. The resulting good 2D classes were selected and corresponding particles were refined with the standard 3D auto-refinement to generate the initial model. This initial model then underwent three runs of CTF refinement in the order of magnification anisotropy, optical aberrations (up to the fourth order) and per-particle defocus and per-micrograph astigmatism. This was followed by a round of 3D auto-refinement before optimizing per-particle beam-induced motion tracks by Bayesian polishing. The polished particles were further subjected to another round of CTF refinement as before. Next, these particles were classified into 3D volumes without particle alignment and the classes containing particles with the highest estimated resolutions were selected for a series of refinements including 3D auto-refinement, CTF refinements and another round of 3D auto-refinement. Finally, the resolution of the reconstructed structure was estimated using the Fourier shell correlation (FSC) criteria of 0.143.

Data processing for Falcon C data was done similarly to the Ceta-F datasets above with modifications to compensate for the EER data format. All datasets were collected on a Falcon C prototype. The resulting output EER files were in a pseudo-Falcon 4i format at half the real pixel size in 4K by 4K so that data could be processed by both CryoSPARC and RELION prior to the release of the 2K EER branch. Data was imported into each of the packages as if it was Falcon 4i data, “EER upsampling = 1” parameter was set for motion correction in CryoSPARC while in RELION, motion correction was run with default values which imposes “EER upsampling = 1” by default. In both packages, motion correction output is binned to 2K by 2K and the rest of the processing is carried out as per the protocol below. The pseudo-4K EER format allows us to benefit from CryoSPARC Live on-the-fly processing and monitoring during data collection and also access the full RELION data processing pipeline before both packages support 2K EER, which is the native image format of Falcon C. From the gallery of 2D class averages generated by CryoSPARC Live, the promising classes of particles were then selected and exported. Particle positions of selected particles were converted via pyEM (Asarnow et al., 2019) for particle extraction in RELION and processed in the manner described in the previous paragraph starting from 2D classification.

Specifically, the CAK dataset contained 5,036 exposures of which 4,275 were accepted after motion correction and CTF fit resolution curation. A three-picking-type protocol, which merges picks from blob picking in CryoSPARC, 2D and 3D template picking output between 2.0 and 3.5 million particles per run, of which around 400,000 to 450,000 survived the first round of 2D classification for each strategy. After the removal of duplicates, a subset of 960,000 particles was subject to another 2D classification, resulting in 833,917 cleaned-up particles, which were imported into RELION. Following several rounds of 3D classification, iterative CTF refinement and particle polishing, the final map was calculated from 181,120 particles using Blush regularisation in RELION 5.0 (Kimanius et al., 2024).

### Map-model FSC calculations

Small errors in the pixel size of maps were corrected by comparison to high-resolution PDB models using the fit in map function in Chimera (Pettersen et al., 2021). The atomic models were automatically and iteratively refined using phenix.real_space_refine using default parameters and electron scattering table (Afonine et al., 2018). Finaly, the refined models were used as inputs for phenix.validation_cryoem to calculate a FSC (map-model). Map-model FSC values were calculated using single monomers and full maps, with the masks calculated from the models. The PDB codes for the initial models used for building were as follows (structure: initial model): T20S proteosome: 1PMA, GABA_A_ receptor: 7A5V, apoferritin: 7A4M, haemoglobin: 5NI1, rabbit muscle aldolase: 6ALD, CDK-activating kinase complex: 8P6Y, transthyretin: 4PVN.

## Data availability

High-resolution cryo-EM maps have been deposited to the Electron Microscopy Data Bank (EMDB) using accession codes EMD-13769 (Apoferritin, Ceta-F), EMD-13816 (GABA_A_, Ceta-F), EMD-14249 (T20S, Ceta-F), EMD-51463 (Apoferritin 0.55 Å/pixel, Falcon C), EMD-51481 (Apoferritin 0.7 Å/pixel, Falcon C), EMD-51483 (Apoferritin 0.9 Å/pixel, Falcon C), EMD-51587 (T20S 0.7 Å/pixel, Falcon C), EMD-51503 (T20S 0.9 Å/pixel, Falcon C), EMD-51506 (GABA_A_ 0.7 Å/pixel, Falcon C), EMD-51508 (GABA_A_ 0.9 Å/pixel, Falcon C), EMD-51511 (Aldolase 0.55 Å/pixel, Falcon C), EMD-51512 (Aldolase 0.7 Å/pixel, Falcon C), EMD-51519 (CAK complex, Falcon C), EMD-51525 (Haemoglobin, Falcon C) and EMD-51526 (TTR, Falcon C). Electron micrograph movies for selected datasets have been deposited to the Electron Microscopy Public Image Archive (EMPIAR) with accession codes EMPIAR-10844 (Apoferritin, Ceta-F), EMPIAR-10858 (GABA_A_, Ceta-F) and EMPIAR-10961 (T20S, Ceta-F).

## Acknowledgements

This work was supported by Thermo Fisher Scientific and carried out on a Tundra microscope equipped with a Ceta-F detector and later exchanged to a Falcon C detector within the Thermo Fisher Scientific RnD facility. We thank Hanna Halabuková and Zuzana Hlavenková for their support with the microscope and logistics. Work on GABA_A_ receptor was supported by the UK Medical Research Council grant MC_UP_1201/15 to A.R.A. and the work on T20S was supported from the Max Planck Society, transthyretin work was supported by the National Institutes of Health (NIH) grant GM14305 to G.C.L. B.J.G. was supported by a career development fellowship from the Medical Research Council of the UK (grant number MR/V009354/1) and V.I.C. was funded by an ICR PhD studentship.

## Author Contributions

D.K., L.Y and A.K. conceived the project. D.K., A.F. K, and W.Y. collected all datasets and processed apoferritin, proteasome T20S, GABAA, Haemoglobin, AAV9 and transthyretin. B.J.G and A.F.K processed CAK dataset. D.K. prepared apoferritin, Haemoglobin and AAV9 samples, D.B.M, S.K, V.I.C and B.B prepared GABAA, T20S, CAK and transthyretin samples respectively. D.K., A.F.K, A.K, D.B.M, G.C.L, A.A.R and B.J.G interpreted the cryo-EM data. O.V., M.M, O. S., and V. D. supported microscope alignments and installation and testing of Ceta-F and Falcon C detectors. D.K., A.F.K, L.Y. and A.K wrote the manuscript with the support from A.A.R., J.P., G.C.L. and B.J.G

## Supplementary Information

**Table S1.** Parameters used for calculating CTF envelope function. *ΔE* is energy spread of the electron source, *E* is the accelerating voltage, *λ* is wavelength of the electron at this accelerating voltage, 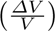 is the relative RMS instability of the high tension, and 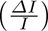 is the relative RMS instability of the objective lens

| | $C_C$ (mm) | $C_C$ (mm) | $\Delta E$ (eV) | $E$ (keV) | $\lambda$ (pm) | $\frac{\Delta V}{V}$ (ppm) | $\frac{\Delta I}{I}$ (ppm) |
| --- | --- | --- | --- | --- | --- | --- | --- |
| Tundra w/ SP-TWIN | 1.7 | 1.6 | 0.8 | 100 | 3.70 | 0.07 | 0.1 |
| Krios w/ C-TWIN | 2.7 | 2.6 | 0.8 | 300 | 1.97 | 0.07 | 0.1 |

**Figure S1.**
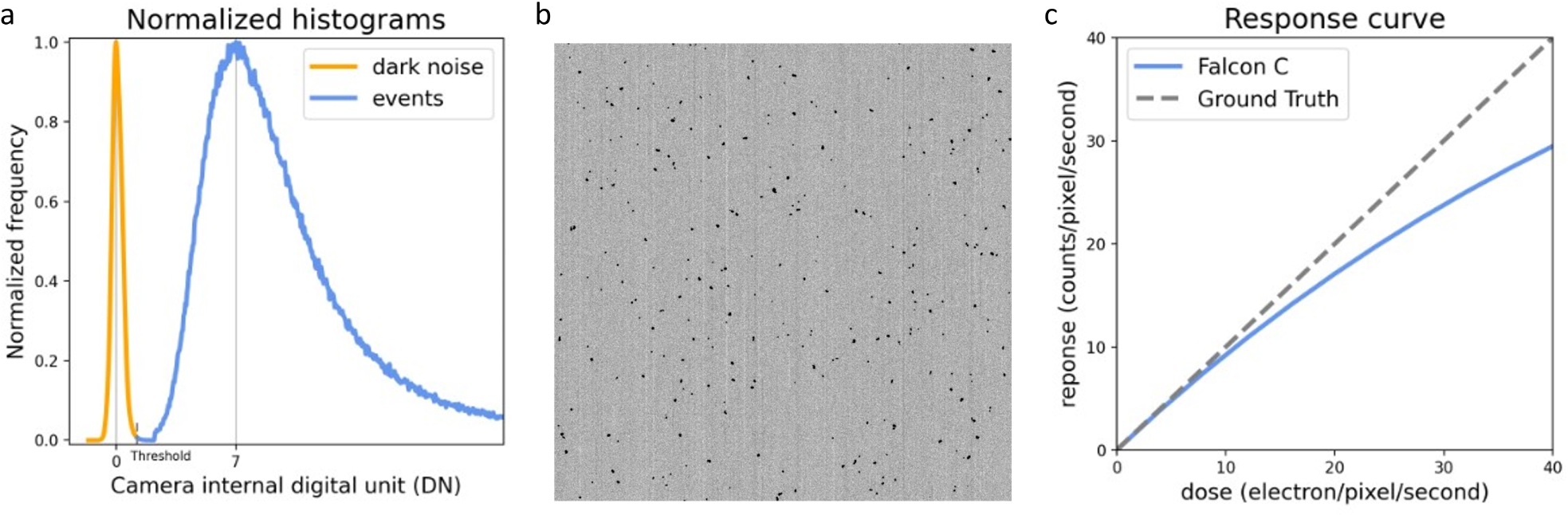
Characterization of Falcon C detector at 100 keV. (**a**) Normalized histogram of noise, Gaussian distributed with a standard deviation of 0.36 DN and electron events, Landau distributed with a mode of 7 DN. (**b**) A 512 x 512 patch of the Falcon C detector readout, input for electron counting. (**c**) Dose-response curve of Falcon C at 100 keV.

**Figure S2.**
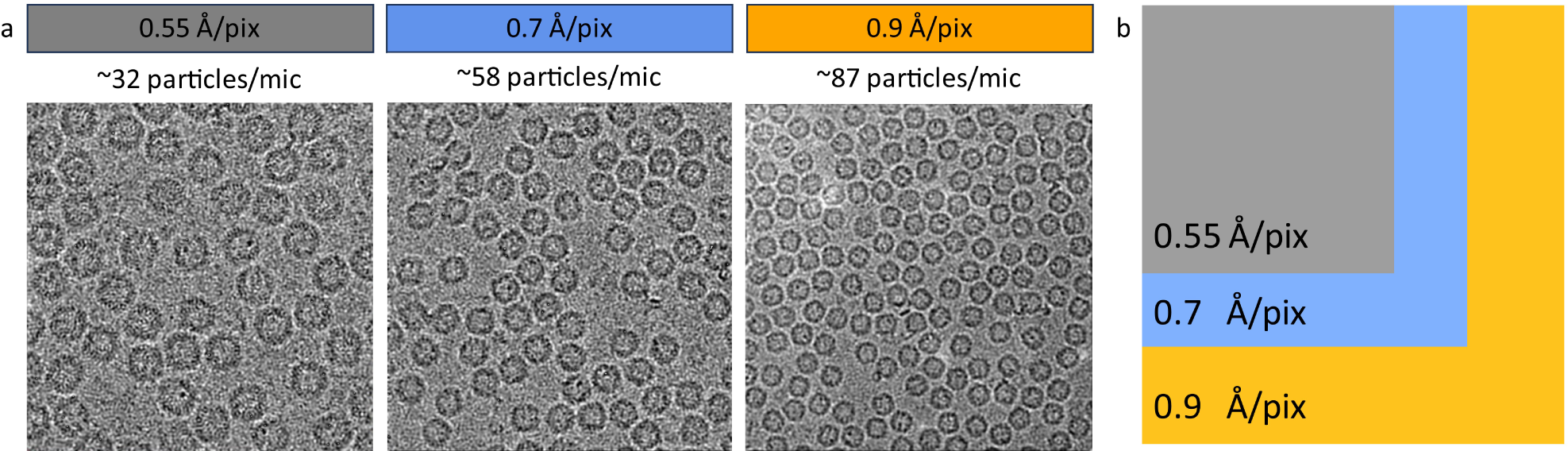
Field of view on Falcon C detector. (**a**) Representative micrographs of the apoferritin sample in vitreous ice imaged at three different magnifications (0.55 Å/pix, 0.7 Å/pix and 0.9 Å/pix) showing the corresponding field of view, including the average number of particles per view. (**b**) Comparison of relative sizes of the field of view at different magnifications.

**Figure S3.**
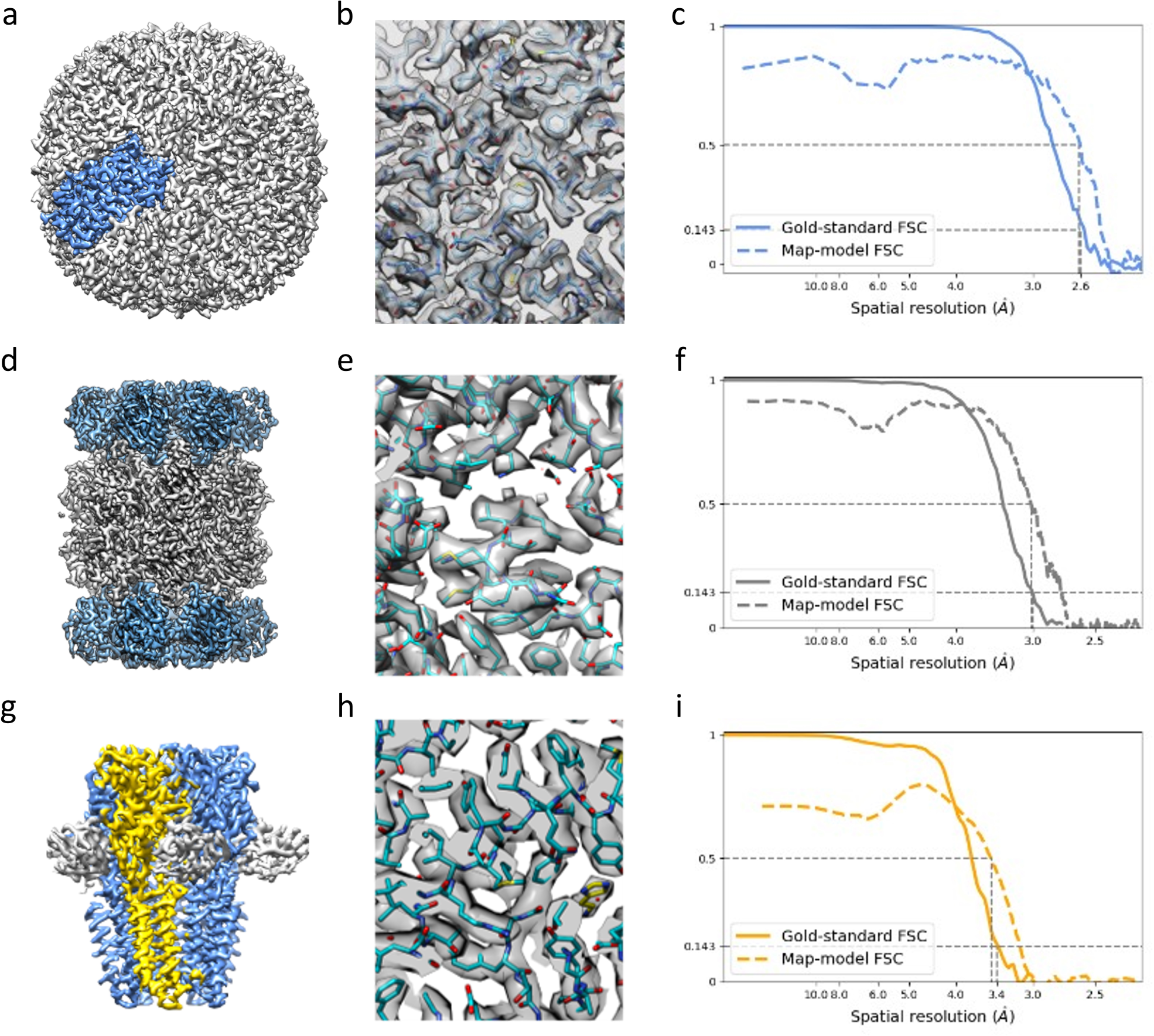
Cryo-EM structures of apoferritin, 20S proteasome and GABA_A_ receptor determined at 100 keV on Ceta-F detector. (**a**) A 2.6 Å reconstruction of apoferritin with octahedral symmetry. The asymmetric unit is highlighted in blue. (**b**) A close-up view of the EM density with the fitted atomic model. (**c**) Gold-standard FSC curve and map-to-model FSC curve corresponding to the map in (a). (**d**) A 3.0 Å reconstruction of proteasome 20S with D7 symmetry. (**e**) A close-up view of the EM density with the fitted atomic model. (**f**) Gold-standard FSC curve and map-to-model FSC curve corresponding to the map in (d). (**g**) A 3.4 Å reconstruction of GABA_A_ receptor with C5 symmetry. (**h**) A close-up view of the EM density with the fitted atomic model. (**i**) Gold-standard FSC curve and map-to-model FSC curve corresponding to the map in (g).

**Figure S4.**
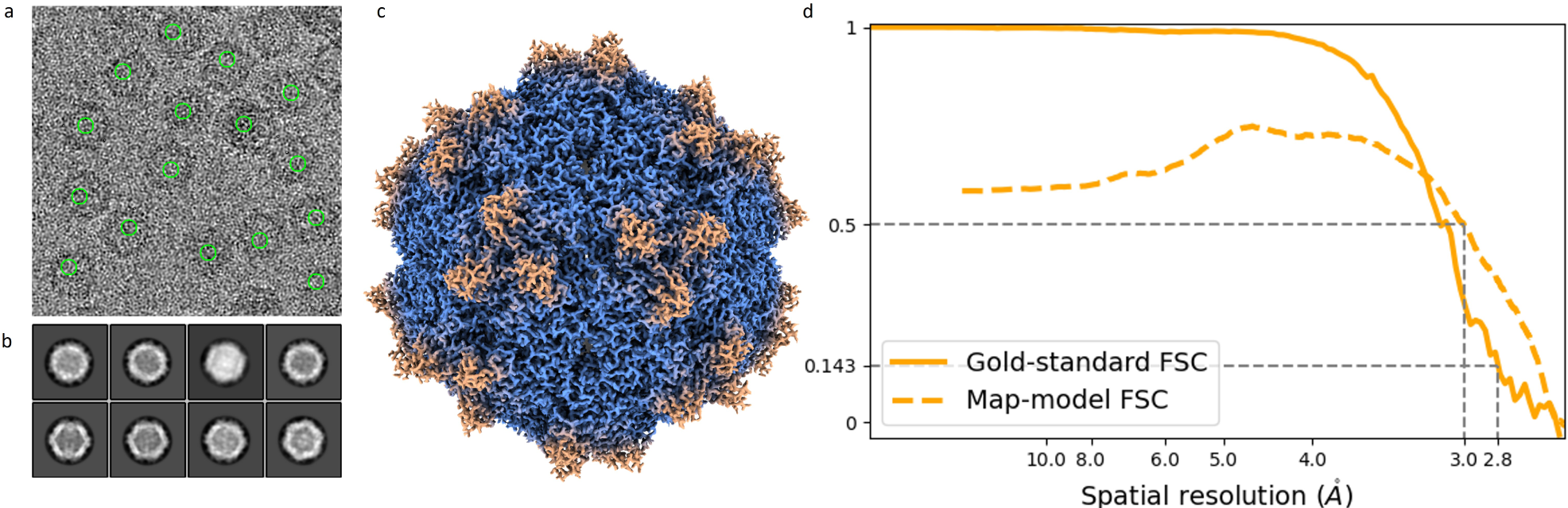
Cryo-EM structures of AAV9 determined at 100 keV on Falcon C detector. (**a**) Representative micrographs of the AAV9 sample in vitreous ice. (**b**) Reference free 2D class averages showing mix population of full, partially filled, and empty capsids. (**c**) A 2.8 Å reconstruction of AAV9 capsid. (**d**) Gold-standard FSC curve and map-to-model FSC curve corresponding to the map in (c).

**Table S2.** Cryo-EM data collection parameters, processing parameters and reconstruction results when using the Ceta-F detector.

|  | Apoferritin<br>(C-TWIN) | Apoferritin<br>(SP-TWIN) | GABA<sub>A</sub><br>receptor | T20S<br>proteasome | Haemoglobin |
| --- | --- | --- | --- | --- | --- |
| Image pixel size (Å/pixel) | 0.73 | 0.75 | 0.75 | 0.75 | 0.57 |
| Nominal magnification | 190kx | 180kx | 180kx | 180kx | 240kx |
| Dose on sample (e<sup>-</sup>/Å<sup>2</sup>) | 40 | 30 | 35 | 35 | 48 |
| Flux on detector (e<sup>-</sup>/pixel/s) | 22 | 17.78 | 14.66 | 14.66 | 31 |
| Exposure time (s) | 0.97 | 1.13 | 1.35 | 1.35 | 0.51 |
| Target defocus range (μm) | 0.2-0.6 | 0.2-0.6 | 0.4-0.7 | 0.3-0.6 | 0.5-1.5 |
| Number of movies | 2,395 | 1,510 | 9,643 | 5,176 | 2,011 |
| Initial particle number | 413,180 | 301,075 | 807,374 | 786,979 | 491,473 |
| Final particle number | 234,568 | 153,488 | 92,997 | 183,486 | 41,791 |
| Symmetry imposed | O | O | C5 | D7 | C2 |
| Map resolution at FSC=0.143 (Å) | 3 | 2.6 | 3.4 | 3 | 8.1 |
| B factor (Å<sup>2</sup>) | 234.4 | 173.8 | 207.7 | 211.8 | - |

**Table S3.** Cryo-EM data collection parameters, processing parameters and reconstruction results when only using 4 hours of data from the Falcon C detector.

|  | Apoferritin | T20S Proteasome | GABA<sub>A</sub> receptor | Aldolase |
| --- | --- | --- | --- | --- |
| Image pixel size (Å/pixel) | 0.7 | 0.7 | 0.7 | 0.7 |
| Nominal magnification | 370kx | 370kx | 370kx | 370kx |
| Dose on sample (e<sup>-</sup>/Å<sup>2</sup>) | 40 | 40 | 40 | 50 |
| Flux on detector (e<sup>-</sup>/pixel/s) | 6.8 | 4.4 | 7.2 | 7.12 |
| Exposure time (s) | 2.87 | 4.52 | 2.77 | 3.49 |
| Target defocus range (µm) | 0.2-0.6 | 0.2-0.6 | 0.3-0.9 | 0.3-0.9 |
| Number of movies | 1,000 | 1,000 | 1,000 | 1,000 |
| Initial particle number | 47,839 | 12,443 | 23,499 | 35,254 |
| Final particle number | 46,275 | 7,721 | 17,526 | 14,675 |
| Symmetry imposed | O | D7 | C5 | D2 |
| Map resolution at FSC=0.143 (Å) | 2.3 | 3.1 | 3.3 | 3.4 |

**Table S4.**
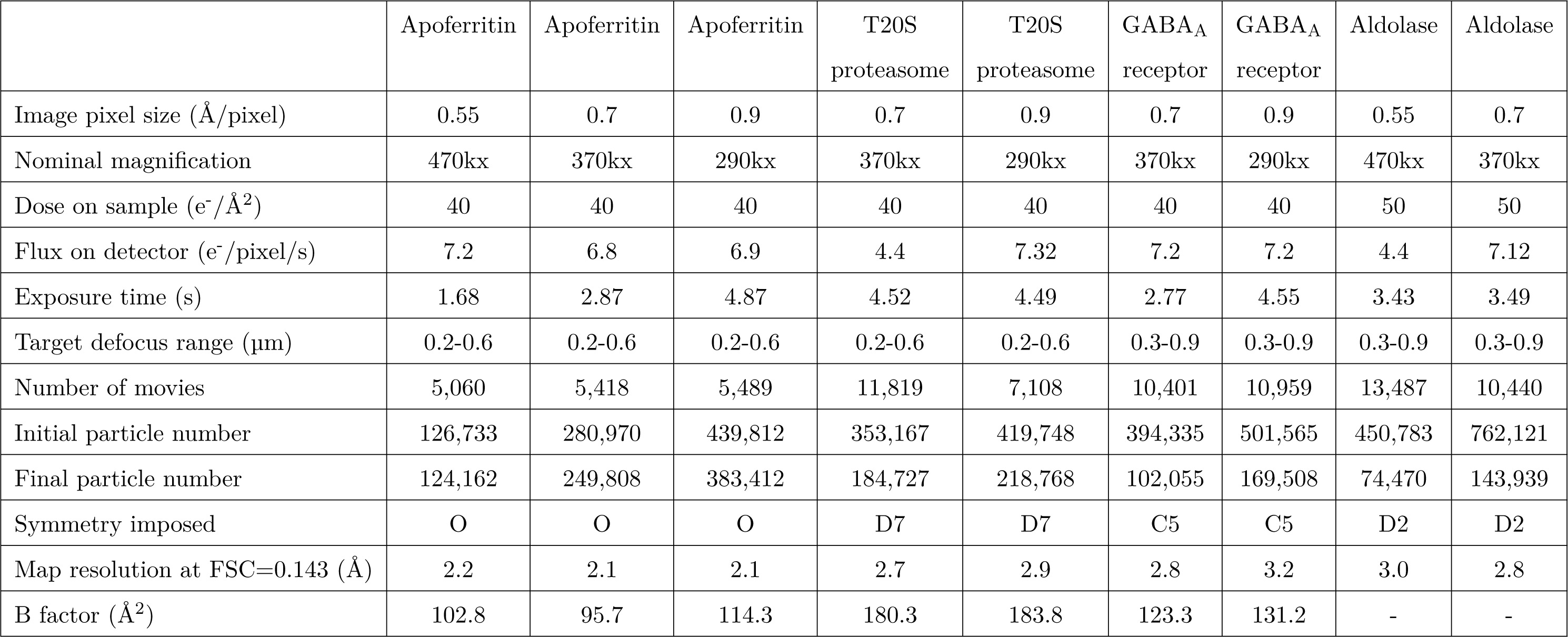
Cryo-EM data collection parameters, processing parameters and reconstruction results when the Falcon C detector with different pixel sizes.

|  | Apoferritin | Apoferritin | Apoferritin | T20S<br>proteasome | T20S<br>proteasome | GABA<sub>A</sub><br>receptor | GABA<sub>A</sub><br>receptor | Aldolase | Aldolase |
| --- | --- | --- | --- | --- | --- | --- | --- | --- | --- |
| Image pixel size (Å/pixel) | 0.55 | 0.7 | 0.9 | 0.7 | 0.9 | 0.7 | 0.9 | 0.55 | 0.7 |
| Nominal magnification | 470kx | 370kx | 290kx | 370kx | 290kx | 370kx | 290kx | 470kx | 370kx |
| Dose on sample (e<sup>-</sup>/Å<sup>2</sup>) | 40 | 40 | 40 | 40 | 40 | 40 | 40 | 50 | 50 |
| Flux on detector (e<sup>-</sup>/pixel/s) | 7.2 | 6.8 | 6.9 | 4.4 | 7.32 | 7.2 | 7.2 | 4.4 | 7.12 |
| Exposure time (s) | 1.68 | 2.87 | 4.87 | 4.52 | 4.49 | 2.77 | 4.55 | 3.43 | 3.49 |
| Target defocus range (µm) | 0.2-0.6 | 0.2-0.6 | 0.2-0.6 | 0.2-0.6 | 0.2-0.6 | 0.3-0.9 | 0.3-0.9 | 0.3-0.9 | 0.3-0.9 |
| Number of movies | 5,060 | 5,418 | 5,489 | 11,819 | 7,108 | 10,401 | 10,959 | 13,487 | 10,440 |
| Initial particle number | 126,733 | 280,970 | 439,812 | 353,167 | 419,748 | 394,335 | 501,565 | 450,783 | 762,121 |
| Final particle number | 124,162 | 249,808 | 383,412 | 184,727 | 218,768 | 102,055 | 169,508 | 74,470 | 143,939 |
| Symmetry imposed | O | O | O | D7 | D7 | C5 | C5 | D2 | D2 |
| Map resolution at FSC=0.143 (Å) | 2.2 | 2.1 | 2.1 | 2.7 | 2.9 | 2.8 | 3.2 | 3.0 | 2.8 |
| B factor (Å<sup>2</sup>) | 102.8 | 95.7 | 114.3 | 180.3 | 183.8 | 123.3 | 131.2 | - | - |

**Table S5.**
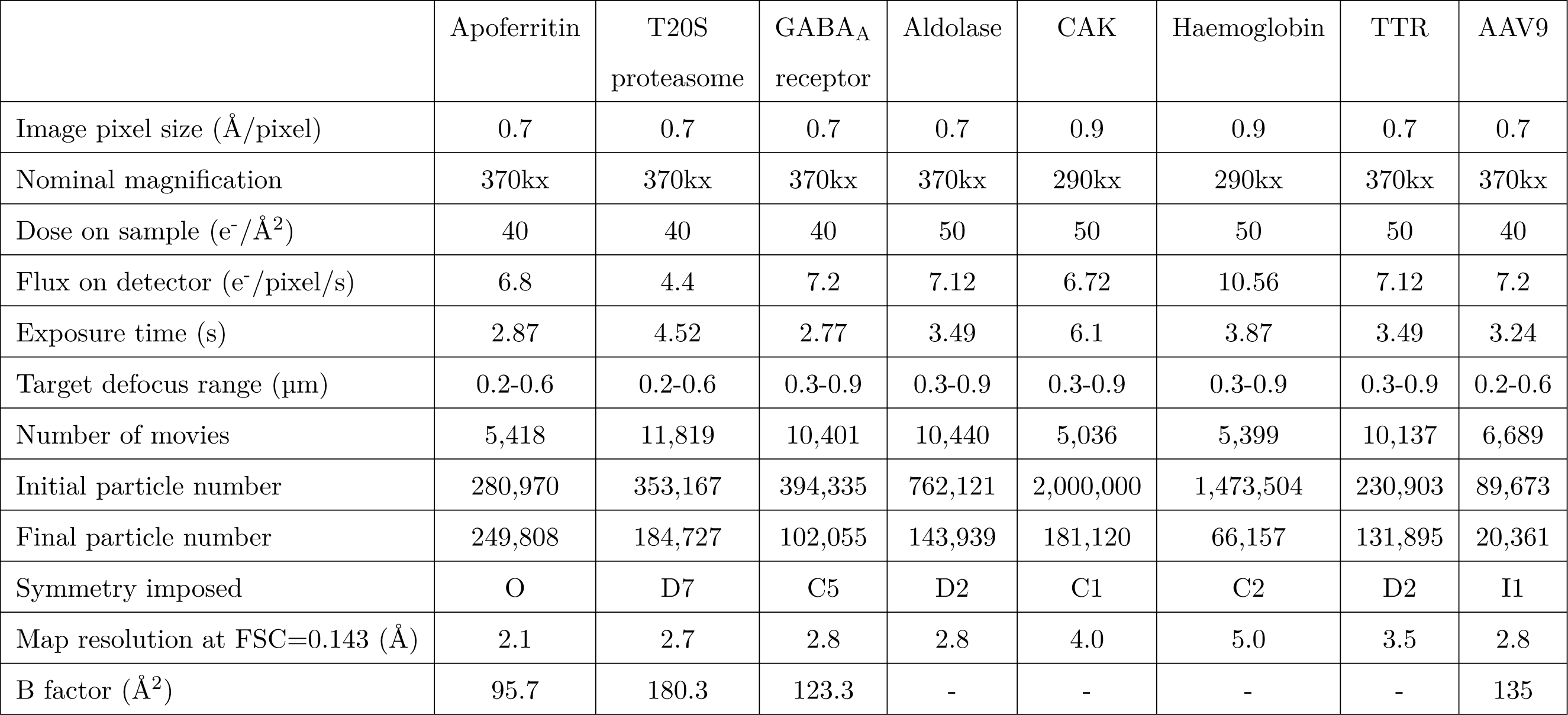
Cryo-EM data collection parameters, processing parameters and the best reconstruction resolution achieved when the Falcon C detector.

|  | Apo ferritin | T20S<br>proteasome | GABA<sub>A</sub><br>receptor | Aldolase | CAK | Haemoglobin | TTR | AAV9 |
| --- | --- | --- | --- | --- | --- | --- | --- | --- |
| Image pixel size (Å/pixel) | 0.7 | 0.7 | 0.7 | 0.7 | 0.9 | 0.9 | 0.7 | 0.7 |
| Nominal magnification | 370kx | 370kx | 370kx | 370kx | 290kx | 290kx | 370kx | 370kx |
| Dose on sample (e<sup>-</sup>/Å<sup>2</sup>) | 40 | 40 | 40 | 50 | 50 | 50 | 50 | 40 |
| Flux on detector (e<sup>-</sup>/pixel/s) | 6.8 | 4.4 | 7.2 | 7.12 | 6.72 | 10.56 | 7.12 | 7.2 |
| Exposure time (s) | 2.87 | 4.52 | 2.77 | 3.49 | 6.1 | 3.87 | 3.49 | 3.24 |
| Target defocus range (µm) | 0.2-0.6 | 0.2-0.6 | 0.3-0.9 | 0.3-0.9 | 0.3-0.9 | 0.3-0.9 | 0.3-0.9 | 0.2-0.6 |
| Number of movies | 5,418 | 11,819 | 10,401 | 10,440 | 5,036 | 5,399 | 10,137 | 6,689 |
| Initial particle number | 280,970 | 353,167 | 394,335 | 762,121 | 2,000,000 | 1,473,504 | 230,903 | 89,673 |
| Final particle number | 249,808 | 184,727 | 102,055 | 143,939 | 181,120 | 66,157 | 131,895 | 20,361 |
| Symmetry imposed | O | D7 | C5 | D2 | C1 | C2 | D2 | I1 |
| Map resolution at FSC=0.143 (Å) | 2.1 | 2.7 | 2.8 | 2.8 | 4.0 | 5.0 | 3.5 | 2.8 |
| B factor (Å<sup>2</sup>) | 95.7 | 180.3 | 123.3 | - | - | - | - | 135 |

## Bibliography

Afonine, P. V., Klaholz, B. P., Moriarty, N. W., Poon, B. K., Sobolev, O. V., Terwilliger, T. C., Adams, P. D., and Urzhumtsev, A. New tools for the analysis and validation of cryo-EM maps and atomic models. Acta Crystallographica Section D: Structural Biology, 74(9):814–840, Sept. 2018. doi: 10.1107/S2059798318009324. Publisher: International Union of Crystallography.

Aricescu, A. R., Lu, W., and Jones, E. Y. A time- and cost-efficient system for high-level protein production in mammalian cells. Acta Crystallographica. Section D, Biological Crystallography, 62(Pt 10):1243–1250, Oct. 2006. doi: 10.1107/S0907444906029799.

Asarnow, D., Palovcak, E., and Cheng, Y. asarnow/pyem: UCSF pyem v0.5, Dec. 2019. URL https://zenodo.org/records/3576630.

Basanta, B., Nugroho, K., Yan, N. L., Kline, G. M., Powers, E. T., Tsai, F. J., Wu, M., Hansel-Harris, A., Chen, J. S., Forli, S., Kelly, J. W., and Lander, G. C. The conformational landscape of human transthyretin revealed by cryo-EM. bioRxiv: The Preprint Server for Biology, page 2024.01.23.576879, Jan. 2024. doi: 10.1101/2024.01.23.576879.

Cheng, A., Eng, E. T., Alink, L., Rice, W. J., Jordan, K. D., Kim, L. Y., Potter, C. S., and Carragher, B. High resolution single particle cryo-electron microscopy using beam-image shift. Journal of Structural Biology, 204(2):270–275, Nov. 2018. doi: 10.1016/j.jsb.2018.07.015.

Cushing, V. I., Koh, A. F., Feng, J., Jurgaityte, K., Bondke, A., Kroll, S. H. B., Barbazanges, M., Scheiper, B., Bahl, A. K., Barrett, A. G. M., Ali, S., Kotecha, A., and Greber, B. J. High-resolution cryo-EM of the human CDK-activating kinase for structure-based drug design. Nature Communications, 15(1):2265, Mar. 2024. doi: 10.1038/s41467-024-46375-9. Publisher: Nature Publishing Group.

Guaita, M., Watters, S. C., and Loerch, S. Recent advances and current trends in cryo-electron microscopy. Current Opinion in Structural Biology, 77:102484, Dec. 2022. doi: 10.1016/j.sbi.2022.102484.

Guo, H., Franken, E., Deng, Y., Benlekbir, S., Singla Lezcano, G., Janssen, B., Yu, L., Ripstein, Z. A., Tan, Y. Z., and Rubinstein, J. L. Electron-event representation data enable efficient cryoEM file storage with full preservation of spatial and temporal resolution. IUCrJ, 7(5):860–869, Sept. 2020. doi: 10.1107/S205225252000929X. Publisher: International Union of Crystallography.

Hamdi, F., Tüting, C., Semchonok, D. A., Visscher, K. M., Kyrilis, F. L., Meister, A., Skalidis, I., Schmidt, L., Parthier, C., Stubbs, M. T., and Kastritis, P. L. 2.7 Å cryo-EM structure of vitrified M. musculus H-chain apoferritin from a compact 200 keV cryo-microscope. PloS One, 15(5):e0232540, 2020. doi: 10.1371/journal.pone.0232540.

Herzik, M. A., Wu, M., and Lander, G. C. Achieving better-than-3-Å resolution by single-particle cryo-EM at 200 keV. Nature Methods, 14(11):1075–1078, Nov. 2017. doi: 10.1038/nmeth.4461.

Herzik, M. A., Wu, M., and Lander, G. C. High-resolution structure determination of sub-100 kDa complexes using conventional cryo-EM. Nature Communications, 10(1):1032, Mar. 2019. doi: 10.1038/s41467-019-08991-8. Publisher: Nature Publishing Group.

Janssen, B. J., Hoften, G. C. V., and Luecken, U. Method for acquiring data with an image sensor, Aug. 2014.

Kimanius, D., Jamali, K., Wilkinson, M. E., Lövestam, S., Velazhahan, V., Nakane, T., and Scheres, S. H. W. Data-driven regularization lowers the size barrier of cryo-EM structure determination. Nature Methods, 21(7):1216–1221, July 2024. doi: 10.1038/s41592-024-02304-8. Publisher: Nature Publishing Group.

Koh, A., Khavnekar, S., Yang, W., Karia, D., Cats, D., van der Ploeg, R., Grollios, F., Raschdorf, O., Kotecha, A., and Němeček, D. Routine Collection of High-Resolution cryo-EM Datasets Using 200 KV Transmission Electron Microscope. Journal of Visualized Experiments: JoVE, (181), Mar. 2022. doi: 10.3791/63519.

Kuijper, M., van Hoften, G., Janssen, B., Geurink, R., De Carlo, S., Vos, M., van Duinen, G., van Haeringen, B., and Storms, M. FEI’s direct electron detector developments: Embarking on a revolution in cryo-TEM. Journal of Structural Biology, 192(2): 179–187, Nov. 2015. doi: 10.1016/j.jsb.2015.09.014.

Kühlbrandt, W. The Resolution Revolution. Science, 343(6178):1443–1444, Mar. 2014. doi: 10.1126/science.1251652. Publisher: American Association for the Advancement of Science.

Laverty, D., Desai, R., Uchański, T., Masiulis, S., Stec, W. J., Malinauskas, T., Zivanov, J., Pardon, E., Steyaert, J., Miller, K. W., and Aricescu, A. R. Cryo-EM structure of the human α1β3γ2 GABAA receptor in a lipid bilayer. Nature, 565(7740):516–520, Jan. 2019. doi: 10.1038/s41586-018-0833-4.

Liao, M., Cao, E., Julius, D., and Cheng, Y. Structure of the TRPV1 ion channel determined by electron cryo-microscopy. Nature, 504(7478):107–112, Dec. 2013. doi: 10.1038/nature12822. Publisher: Nature Publishing Group.

Liz, M. A., Coelho, T., Bellotti, V., Fernandez-Arias, M. I., Mallaina, P., and Obici, L. A Narrative Review of the Role of Transthyretin in Health and Disease. Neurology and Therapy, 9(2):395–402, Dec. 2020. doi: 10.1007/s40120-020-00217-0.

Malínský, M., Hoften, G. v., Vyroubal, O., Doležal, V., Č ervinková, M., and Yu, L. CETA-F: Scintillator camera for Entry level 100kV Single Particle Analysis. Microscopy and Microanalysis, 27(S1):1640–1641, Aug. 2021. doi: 10.1017/S1431927621006048.

Masiulis, S., Desai, R., Uchański, T., Serna Martin, I., Laverty, D., Karia, D., Malinauskas, T., Zivanov, J., Pardon, E., Kotecha, A., Steyaert, J., Miller, K. W., and Aricescu, A. R. GABAA receptor signalling mechanisms revealed by structural pharmacology. Nature, 565(7740):454–459, Jan. 2019. doi: 10.1038/s41586-018-0832-5.

McMullan, G., Faruqi, A. R., and Henderson, R. Chapter One - Direct Electron Detectors. In Crowther, R. A., editor, *Methods in Enzymology*, volume 579 of *The Resolution Revolution: Recent Advances In cryoEM*, pages 1–17. Academic Press, Jan. 2016. doi: 10.1016/bs.mie.2016.05.056.

McMullan, G., Naydenova, K., Mihaylov, D., Yamashita, K., Peet, M. J., Wilson, H., Dickerson, J. L., Chen, S., Cannone, G., Lee, Y., Hutchings, K. A., Gittins, O., Sobhy, M. A., Wells, T., El-Gomati, M. M., Dalby, J., Meffert, M., Schulze-Briese, C., Henderson, R., and Russo, C. J. Structure determination by cryoEM at 100 keV. Proceedings of the National Academy of Sciences, 120 (49):e2312905120, Dec. 2023. doi: 10.1073/pnas.2312905120. Publisher: Proceedings of the National Academy of Sciences.

Miller, P. S. and Aricescu, A. R. Crystal structure of a human GABAA receptor. Nature, 512(7514):270–275, Aug. 2014. doi: 10.1038/nature13293. Publisher: Nature Publishing Group.

Nakane, T., Kotecha, A., Sente, A., McMullan, G., Masiulis, S., Brown, P. M. G. E., Grigoras, I. T., Malinauskaite, L., Malinauskas, T., Miehling, J., Uchański, T., Yu, L., Karia, D., Pechnikova, E. V., de Jong, E., Keizer, J., Bischoff, M., McCormack, J., Tiemeijer, P., Hardwick, S. W., Chirgadze, D. Y., Murshudov, G., Aricescu, A. R., and Scheres, S. H. W. Single-particle cryo-EM at atomic resolution. Nature, 587(7832):152–156, Nov. 2020. doi: 10.1038/s41586-020-2829-0.

Naydenova, K., McMullan, G., Peet, M. J., Lee, Y., Edwards, P. C., Chen, S., Leahy, E., Scotcher, S., Henderson, R., and Russo, C. J. CryoEM at 100 keV: a demonstration and prospects. IUCrJ, 6(Pt 6):1086–1098, Nov. 2019. doi: 10.1107/S2052252519012612.

Peet, M. J., Henderson, R., and Russo, C. J. The energy dependence of contrast and damage in electron cryomicroscopy of biological molecules. Ultramicroscopy, 203:125–131, Aug. 2019. doi: 10.1016/j.ultramic.2019.02.007.

Pettersen, E. F., Goddard, T. D., Huang, C. C., Meng, E. C., Couch, G. S., Croll, T. I., Morris, J. H., and Ferrin, T. E. UCSF ChimeraX: Structure visualization for researchers, educators, and developers. Protein Science: A Publication of the Protein Society, 30(1):70–82, Jan. 2021. doi: 10.1002/pro.3943.

Punjani, A., Rubinstein, J. L., Fleet, D. J., and Brubaker, M. A. cryoSPARC: algorithms for rapid unsupervised cryo-EM structure determination. Nature Methods, 14(3):290–296, Mar. 2017. doi: 10.1038/nmeth.4169.

Rohou, A. and Grigorieff, N. CTFFIND4: Fast and accurate defocus estimation from electron micrographs. Journal of Structural Biology, 192(2):216–221, Nov. 2015. doi: 10.1016/j.jsb.2015.08.008.

Rosenthal, P. B. and Henderson, R. Optimal determination of particle orientation, absolute hand, and contrast loss in single-particle electron cryomicroscopy. Journal of Molecular Biology, 333(4):721–745, Oct. 2003. doi: 10.1016/j.jmb.2003. 07.013.

Russo, C. J. and Passmore, L. A. Ultrastable gold substrates: Properties of a support for high-resolution electron cryomicroscopy of biological specimens. Journal of Structural Biology, 193(1):33–44, Jan. 2016. doi: 10.1016/j.jsb.2015.11.006.

Sanguinetti, C., Minniti, M., Susini, V., Caponi, L., Panichella, G., Castiglione, V., Aimo, A., Emdin, M., Vergaro, G., and Franzini, M. The Journey of Human Transthyretin: Synthesis, Structure Stability, and Catabolism. Biomedicines, 10(8):1906, Aug. 2022. doi: 10.3390/biomedicines10081906. Number: 8 Publisher: Multidisciplinary Digital Publishing Institute.

Sieghart, W. and Savić, M. M. International Union of Basic and Clinical Pharmacology. CVI: GABAA Receptor Subtype- and Function-selective Ligands: Key Issues in Translation to Humans. Pharmacological Reviews, 70(4):836–878, Oct. 2018. doi: 10.1124/pr.117.014449.

Thangaratnarajah, C., Rheinberger, J., and Paulino, C. Cryo-EM studies of membrane proteins at 200 keV. Current Opinion in Structural Biology, 76:102440, Oct. 2022. doi: 10.1016/j.sbi.2022.102440.

Tiemeijer, P. C., Bischoff, M., Freitag, B., and Kisielowski, C. Using a monochromator to improve the resolution in TEM to below 0.5 Å. Part I: Creating highly coherent monochromated illumination. Ultramicroscopy, 114:72–81, Mar. 2012. doi: 10.1016/j.ultramic.2012.01.008.

Uchański, T., Masiulis, S., Fischer, B., Kalichuk, V., López-Sánchez, U., Zarkadas, E., Weckener, M., Sente, A., Ward, P., Wohlkönig, A., Zögg, T., Remaut, H., Naismith, J. H., Nury, H., Vranken, W., Aricescu, A. R., Pardon, E., and Steyaert, J. Megabodies expand the nanobody toolkit for protein structure determination by single-particle cryo-EM. Nature Methods, 18 (1):60–68, Jan. 2021. doi: 10.1038/s41592-020-01001-6.

Vinothkumar, K. R. and Henderson, R. Single particle electron cryomicroscopy: trends, issues and future perspective. Quarterly Reviews of Biophysics, 49:e13, Jan. 2016. doi: 10.1017/S0033583516000068.

Wu, M., Lander, G. C., and Herzik, M. A. Sub-2 Angstrom resolution structure determination using single-particle cryo-EM at 200keV. Journal of Structural Biology: X, 4:100020, Jan. 2020. doi: 10.1016/j.yjsbx.2020.100020.

Zheng, S. Q., Palovcak, E., Armache, J.-P., Verba, K. A., Cheng, Y., and Agard, D. A. MotionCor2: anisotropic correction of beam-induced motion for improved cryo-electron microscopy. Nature Methods, 14(4):331–332, Apr. 2017. doi: 10.1038/nmeth.4193. Publisher: Nature Publishing Group.

Zivanov, J., Nakane, T., Forsberg, B. O., Kimanius, D., Hagen, W. J., Lindahl, E., and Scheres, S. H. New tools for automated high-resolution cryo-EM structure determination in RELION-3. eLife, 7:e42166, Nov. 2018. doi: 10.7554/eLife.42166. Publisher: eLife Sciences Publications, Ltd.

Zivanov, J., Otón, J., Ke, Z., von Kügelgen, A., Pyle, E., Qu, K., Morado, D., Castaño-Díez, D., Zanetti, G., Bharat, T. A. M., Briggs, J. A. G., and Scheres, S. H. W. A Bayesian approach to single-particle electron cryo-tomography in RELION-4.0. eLife, 11: e83724, Dec. 2022. doi: 10.7554/eLife.83724.

Zwickl, P., Lottspeich, F., and Baumeister, W. Expression of functional Thermoplasma acidophilum proteasomes in Escherichia coli. FEBS letters, 312(2-3):157–160, Nov. 1992. doi: 10.1016/0014-5793(92)80925-7.

